# Bayesian *Occam’s Razor* to Optimize Metamodeling for Complex Biological Systems

**DOI:** 10.1101/2024.05.28.594654

**Authors:** Chenxi Wang, Jihui Zhao, Jingjing Zheng, Barak Raveh, Xuming He, Liping Sun

## Abstract

Modeling complex biological systems necessitates the integration of vast and multifaceted information spanning various aspects of these systems, and is expected to yield more insights into the system than any of the inputs. Metamodeling, a specialized form of integrative modeling, addresses this by integrating existing models. Developing and optimizing metamodels pose challenges due to the complexities introduced by diverse input models and their inherent uncertainties. In this study, we employ Bayesian formalism to rigorously analyze the propagation of probability throughout the metamodeling process and propose quantitative assessments for it. Building on this, we introduce a method for optimizing metamodeling that adheres to the Bayesian *Occam’s razor* rationale, by (i) minimizing model uncertainty; (ii) maximizing model consistency; and (iii) reducing model complexity. To illustrate the benefits of this method, we apply it to the dynamic system of glucose-stimulated insulin secretion in pancreatic *β*-cells. The optimized metamodel delivers more accurate estimates of impaired *β*-cell dynamics and function in T2D subjects compared to the non-optimized one, underscoring the critical role of optimization in enhancing both model reliability and applicability. This method is implemented through the *Integrative Modeling Platform* (IMP), facilitating the development of accurate, precise, and sufficiently simple models for a variety of complex systems.

## Introduction

Models are essential for understanding, modulating, and designing future studies on systems of interest. They are built upon input information (*eg*, physical theories, statistical inference, experimental data, and prior models), and are used to rationalize inputs and make predictions. Within the Bayesian framework, a model can be viewed as a joint distribution over all model variables (or as a Dirac delta function for deterministic models) [1, 2], while also incorporating the statistical relationships among these variables. The modeling process generally involves three steps: (i) define the model representation that specifies the degrees of freedom whose values are determined by modeling, (ii) define a scoring function for ranking alternative models by their agreements with the input information, and (iii) generate an ensemble of good-scoring models through sampling [3, 4]. Ideally, we want to find all models that satisfy the input information, thereby capturing uncertainties in the input information.

Optimization of a model often involves quantitative assessments of how well model variables align with input information under uncertainty throughout the modeling process [5, 6]. In a general modeling process, the representation of a model can be optimized at specific levels of precision (*ie*, coarse-graining level) to facilitate exhaustive sampling [7]. The scoring function can be optimized by, for example, translating sparse Nuclear Magnetic Resonance (NMR) data into likelihoods, effectively capturing their uncertainties in protein structure modeling [8]. The sampling process can be optimized through approaches such as bagging, boosting, and stacking, enabling the generation of model ensembles and the evaluation of model uncertainty [9, 10]. Optimization techniques have significantly advanced diverse modeling tasks in biology, including reconstructing gene regulatory networks from single-cell RNA-Seq [11], predicting metabolic fluxes through metabolic flux balance analysis [12], and improving the accuracy of protein protein docking [13]. Once an optimal model is found and well validated, it can be used to interpret results and formulate testable predictions.

Modeling complex biological systems often necessitates an integrative modeling approach [14, 15]. One example is cell modeling, where a variety of models have been constructed by integrating multiple types of information describing various aspects of the cell. These models include spatial-temporal models, ordinary differential equation models, and hierarchical structural models [16, 17, 18]. Metamodeling, a specialized form of integrative modeling, integrates a collection of input models into a metamodel (*ie*, a model of these models) [19, 20]. From a Bayesian perspective, integrative modeling aims to compute a posterior model density through likelihoods and priors, with a modeling process that includes representation, scoring, and sampling [21]. Similarly, metamodeling proceeds through three stages: (i) convert input models into probabilistic surrogate models; (ii) couple surrogate models; and (iii) update surrogate models and input models *via* backpropagation. Metamodeling has found extensive applications in modeling complex biological systems, including protein structures [15], gene regulatory networks [22], and entire cells [1]. Optimization of metamodeling, however, presents significant challenges due to the complexities introduced by diverse input models and various sources of uncertainty.

Fortunately, Bayesian methods provide a robust probabilistic framework to evaluate model variables under uncertainty. Bayesian probability theory further supports model selection in a manner that embodies *Occam’s Razor* [23, 24], favoring simpler models over complex ones when they demonstrate similar predictive capabilities [25, 26]. In this work, we address the optimization problem through the intricate case of metamodeling for complex biological systems, utilizing the previously established general metamodeling framework based on Probabilistic Graphical Models (PGMs) [27, 1]. We apply it to the dynamics system of glucose-stimulated insulin secretion dynamics in pancreatic *β*-cells, yielding more accurate estimates for impaired *β*-cell dynamics and function in T2D subjects.

## Methods

Model variables can be classified into two distinct types: (i) target variables, which are of primary interest and inform the answers to questions posed by a model; and (ii) nuisance variables, which define the modeling process. The propagation of probability through the modeling process determines the posterior model density of all values of the target variables, given nuisance variables and input information (SI Text 1). The modeling process encompasses representation, scoring, and sampling in general modeling, while, by analogy, it includes conversion, coupling, and backpropagation in metamodeling [28, 21]. We begin with a theoretical foundation for computing the target posterior model density based on input information in a general modeling process (SI Text 2), and, in greater detail, for computing the target posterior model density based on input models at each stage of metamodeling (SI Text 3). We now analytically establish definitive rules for the probability propagation for target variables across metamodeling stages and propose quantitative assessments for it.

### Probability propagation in the conversion stage

First, each input model is converted into its corresponding probabilistic surrogate model. The surrogate model is a potentially simplified input model. It is defined formally by a PGM, which provides both the statistical dependencies between the surrogate model variables (*ie*, surrogate PGM graph) and the joint distribution (*ie*, surrogate PGM distribution) (Fig. 1, Surrogate models) [27]. The conversion involves two steps: (i) select surrogate model variables, often a subset of input model variables and potential additional variables, and (ii) construct the surrogate PGM through various conversion methods such as linear regression fitting [1], Kriging modeling [29], and machine learning methods [30]. The conversion stage is analogous to defining a unified model representation given multiple types of input information, thus specifying the degrees of freedom for the subsequent modeling process. Note that input data can be treated as single-variable models, and can thus be converted into single-variable surrogate models with the same probability distribution.

**Fig 1.**
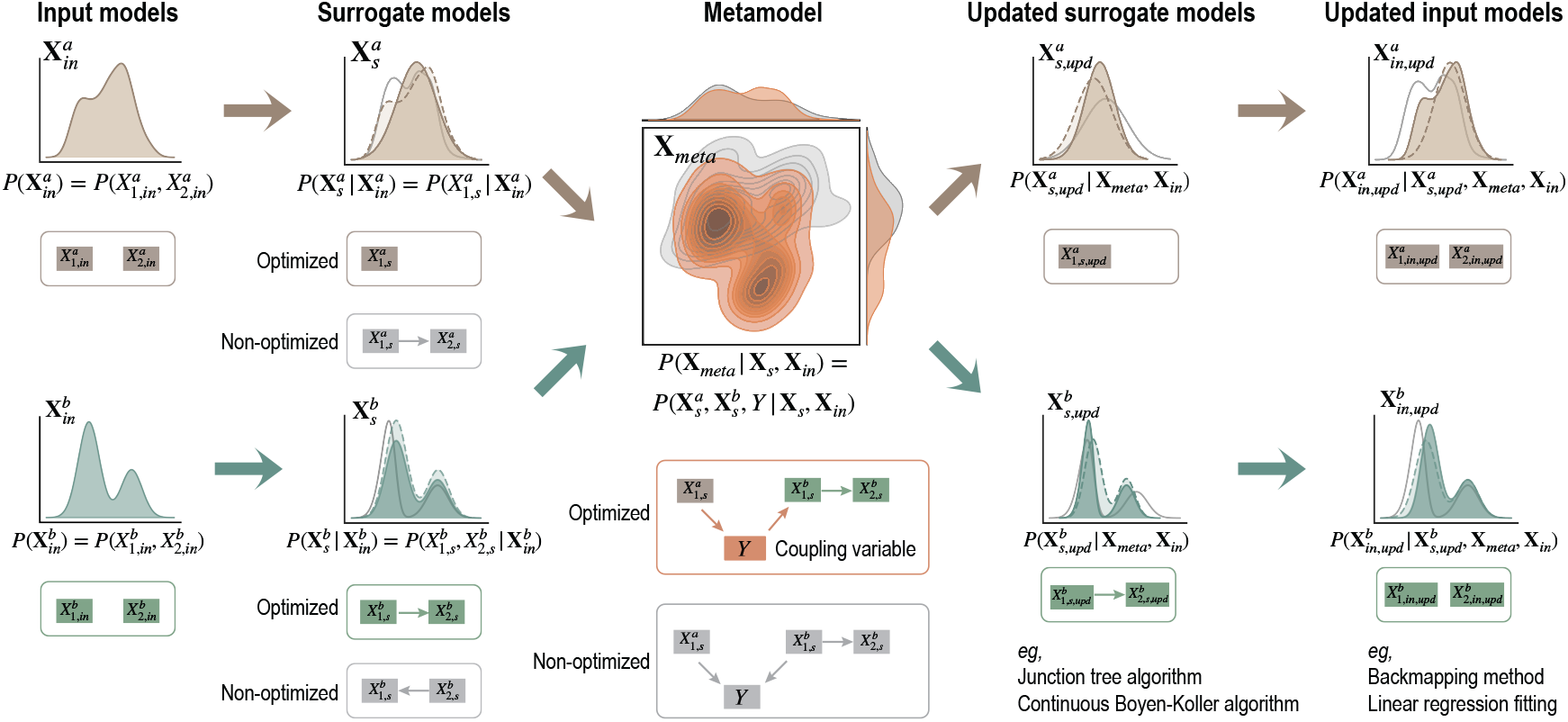
Overview of probability propagation in metamodeling. Input models *a* (khaki) with its distribution 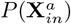 and *b* (green) with its distribution 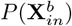,each consisting of two variables. Surrogate model *a* comprises one variable, with its distribution 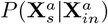;surrogate model *b* comprises two variables, with its distribution 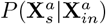.The metamodel constructed using the head-to-tail coupling PGM graph, with its distribution 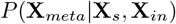.Updated surrogate models *a* and *b* obtained using different algorithms for model inference during backpropagation, with their distributions 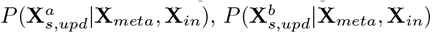, respectively. Updated input models *a* and *b* obtained using different methods during backpropagation, with, their distributions 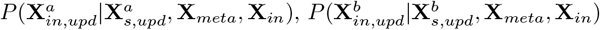, respectively. Thick arrows indicate the propagation of uncertainty across metamodeling stages. Colored solid and dashed lines depict the distributions of optimized models between adjacent stages, while grey solid lines depict distributions of non-optimized ones in each stage. Colored model PGM graphs represent optimized models, while grey ones represent non-optimized ones.

Next, we analytically propagate the probability of a Gaussian input model to its surrogate model. The input model is defined as:

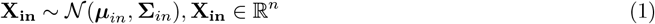

where ***µ***_*in*_ and **Σ**_*in*_ are the mean and covariance matrix of the input model distribution. For example, the input model *a* is a joint distribution 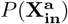 over a set of two random variables 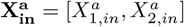 (Fig. 1, Input models). The probability propagation from the input model to the surrogate model follows the chain rule:

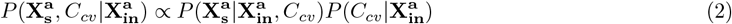

where 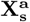 represents surrogate model variables, *C*_*cv*_ is the nuisance variable indicating the conversion method.

The surrogate model distribution over all conceivable conversion methods 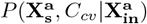 is proportional to the surrogate model distribution given the input model and a single conversion method 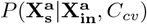,and the distribution of all conversion methods given the input model 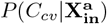. If the conversion method is fixedby the modeler as 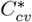 (*ie*, 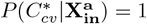), then the surrogate PGM distribution is the joint distribution over its surrogate model variables given the input model, 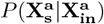.

With the probability distributions of input models and surrogate models in hand, we can quantify the probability propagation in this stage by (i) changes in model uncertainty as the difference in entropy between the input model 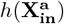 and the surrogate model 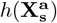;and (ii) overlaps between the marginal distributions of aligned variables, the variables found in both the input and surrogate models [31]. The overlap index for a specific variables pair 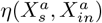 is:

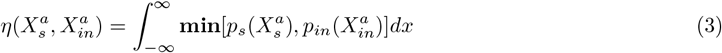

where 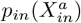 and 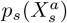 are the marginal distributions (*ie*, probabilistic density functions) of single input and surrogate model variables, respectively. The average of 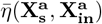,defined as model consistency, reflects the effectiveness of the surrogate models in reproducing the input model. Comparing changes in the uncertainty of surrogate models and consistency between input models and surrogate models can potentially guide us to optimize the conversion stage and select the best surrogate model. Notably, when two candidate surrogate models show comparable uncertainty and consistency with the input model, the optimal surrogate model should be the one with lower complexity (brown PGM in Fig. 1, Surrogate models, consisting of one node and zero edges) compared to the other one (grey PGM in Fig. 1, Surrogate models, consisting of two nodes and one edge).

### Probability propagation in the coupling stage

After converting input models into surrogate models, we couple these surrogate models into a metamodel. The metamodel is defined by a meta PGM, including a meta PGM graph over metamodel variables (*ie*, all surrogate model variables and coupling variables) and a meta PGM distribution spanned over all variables in the meta PGM graph (Fig. 1, Metamodel). The coupling stage is achieved by the construction of a coupling PGM graph through three steps: (i) identify connecting variables across surrogate models; (ii) introduce one or more explicit coupling variables to formally relate the connecting variables; and (iii) specify conditional probability distributions (CPDs) over the connecting variables and the coupling variable(s). In addition, one might construct the coupling PGM graph by defining CPDs between the connecting variables without introducing explicit coupling variables. For example, two subunits in a protein complex can be coupled directly *via* the binding free energy. The coupling stage is analogous to scoring different pieces of information in other model approaches, such as defining a scoring function combining distance and angle restraints in protein modeling [32], or developing a constraint to integrate macroscopic traffic flow models with microscopic agent-based models [33].

Next, we analytically propagate the probability of two static surrogate models to the metamodel. Assuming the surrogate models are constructed using a single conversion method, the probability propagation from the surrogate models to the metamodel follows the chain rule:

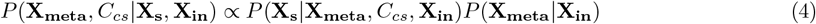

where **X**_**meta**_ represents metamodel variables, *C*_*cs*_ is a nuisance variable indicating the coupling strategy. The meta PGM distribution over all coupling strategies *P* (**X**_**meta**_, *C*_*cs*_ |**X**_**s**_, **X**_**in**_) is proportional to the likelihood *P* (**X**_**s**_ |**X**_**meta**_, *C*_*cs*_, **X**_**in**_) and the prior *P* (**X**_**meta**_ |**X**_**in**_). The likelihood measures how each surrogate model is satisfied by a metamodel given the coupling strategies and input models. The prior reflects prior knowledge about the statistical dependencies between variables across input models.

Suppose we identify connecting variables from two surrogate models with a simple linear relationship 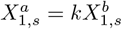,where *k* can be specified by prior knowledge or statistical analysis. We can then introduce an explicit coupling variable *Y*, whose prior distribution *P* (*Y*) is a weighted sum of the marginal distributions of the two connecting variables:

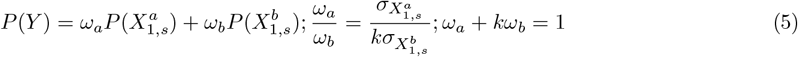

For such a coupling PGM graph composed of two connecting variables and one coupling variable, there are three possible graph structures: head-to-tail, head-to-head, and tail-to-tail. Note that information about any of the nodes along the PGM chain updates the information of other nodes bidirectionally [27, 34]. We now consider the head-to-tail coupling PGM graph, 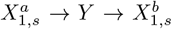. Forwardly, a new estimation for the probability distribution of 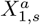 updates 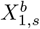 by propagating information over *Y*. Backwardly, observing *Y* updates the estimation of 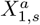.We then specify two Gaussian CPDs for 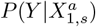 and 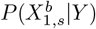 to define the probability propagation along the coupling PGM graph and consequently the coupling strength between two surrogate models:

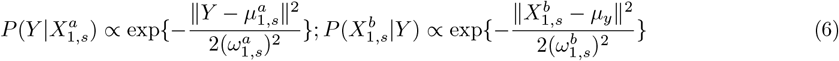

where 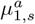 is the mean of the marginal distributions of the connecting variables 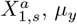, is the mean of the prior distribution of the coupling variable, 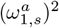 and 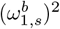 are the variances of the marginal distributions of the connecting variables 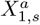 and 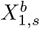, respectively. Note that additional information can be used to inform the prior distribution of the coupling variable *P* (*Y*) (*eg*, assign an observation based on external experiments) and the CPDs (*eg*, assign confidence to each surrogate model based on their uncertainties).

In cases where the modeler fixes a single coupling strategy 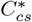 as exemplified here (*ie*, 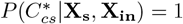,the meta PGM distribution is then the joint distribution over all metamodel variables, given the surrogate and input model variables, *P* (**X**_**meta**_|**X**_**s**_, **X**_**in**_) (Fig. 1, Metamodel). It can be factorized as:

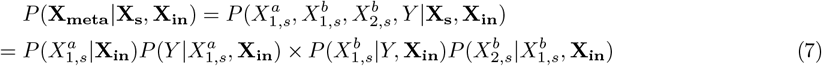

The probability propagation in this stage can be quantified by (i) comparing the difference in surrogate model entropy before and after coupling, and (ii) examining the average overlap index between the marginal distributions of surrogate model variables before and after coupling, indicated by

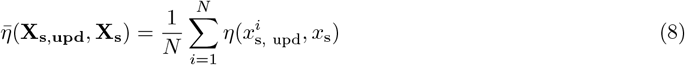

where *N* indicates the number of surrogate model variables. This conveys model consistency and thus reflects the rationalization among pieces of inputs [35]. Comparing changes in the uncertainty of surrogate models and the consistency among them before and after metamodeling can potentially guide us to optimize the coupling stage and select the best coupling PGM graph. For example, the optimal one should yield a metamodel with lower entropy and higher consistency (*eg*, orange PGM in Fig. 1, Metamodel, head-to-tail) compared to the alternative one (*eg*, grey PGM in Fig. 1, Metamodel, head-to-head).

### Probability propagation in the backpropagation stage

After coupling, surrogate models and the corresponding input models are updated by backpropagation. The backpropagation stage is comparable to the process of sampling good-scoring models in a broader modeling context [27]. First, we update the surrogate PGM distributions given the meta PGM distribution obtained in the coupling stage (Fig. 1, Updated surrogate models). Depending on the specific query, this is achieved by either conditioning on or marginalizing out variables in the metamodel using different model inference methods (*eg*, junction tree algorithm[36], continuous Boyen-Koller algorithm [37], reject sampling [38], importance sampling [39]). The updated PGM distribution of each surrogate model Eq.3 is:

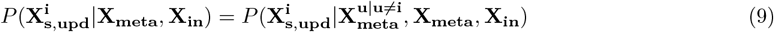

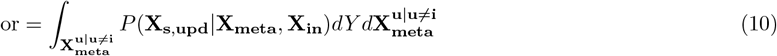

where 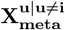 are the variables in the metamodel but not in the surrogate model *i*. Thus, the surrogate model is updated under the contextualization of all other surrogate models. We can also compute the updated distributions spanned by any subsets of variables in the metamodel. Comparing the efficiencies of different inference algorithms facilitates the optimization of the backpropagation stage, which is particularly important when modeling complex systems. We observed a significantly higher efficiency with the junction tree algorithm compared to the continuous Boyen-Koller algorithm (SI Fig. S5). As a result, we have adopted it for all subsequent model inference analyses. Second, depending on the modeler’s interest, one may update the input model based on the updated surrogate model (Fig. 1, Updated input models). For example, the input model of the atomistic protein structure can be updated from the updated surrogate model of the coarse-grained structure using backmapping methods [40]; the input model of a system of ODEs can be updated by linear regression of the time series data sampled from the updated surrogate model [41, 42, 43]. The reconstruction of the updated input model is a separate modeling problem and thus not addressed here.

Next, we compute the updated distribution for surrogate model 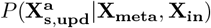 from the meta PGM distribution of a head-to-tail coupling PGM graph 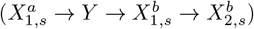 using exact inference [44]:

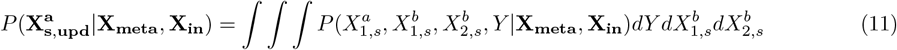

The updated surrogate model is a marginal distribution derived from the joint Gaussian distribution of the metamodel, and thus a joint Gaussian distribution. To exemplify, we compute the mean 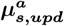 and variance 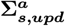 of the updated PGM distribution of surrogate model *a*, which is a single variable model:

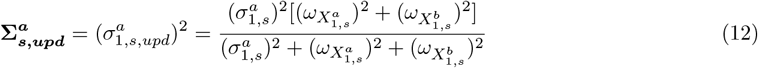

We can thus compute changes in its model uncertainty before and after metamodeling using the ratio of model variances:

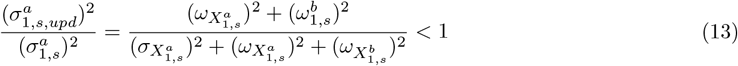

Till here, we have analytically demonstrated that the uncertainty of the updated surrogate model *a* is guaranteed to decrease after metamodeling when employing the head-to-tail coupling PGM graph (see *SI Text 4* for the complete probability calculus). We have provided analytical insights into the propagation of probability across the stages of metamodeling and proposed quantitative assessments for each stage. These analyses guide our optimization of models for complex systems by systematically sampling each stage and comparing model uncertainty, consistency, and complexity.

## Results

We develop a metamodel for glucose-stimulated insulin secretion (GSIS) in pancreatic *β*-cells using three time-dependent Gaussian input models: a system of ordinary differential equations (ODEs) describing the postprandial response (PR) ; a linear model of a pancreatic cell population (Pa) ; and a coarse-grained Brownian dynamics simulation of insulin vesicle exocytosis (VE) [45, 46, 1] (SI Text 5, Table S1-S3, Fig. S1). We optimize the GSIS metamodel by: (i) improving the selection of the surrogate model in the conversion stage, improving the selection of the coupling PGM graph in the coupling stage with and without references, including observations of model variables as an additional data model to resolve conflicts. Finally, we apply the optimized metamodel to reveal *β*-cell dynamics and function in normal and type 2 diabetes (T2D) subjects. We utilize the marginal distributions of aligned variables to quantify probability propagation, rather than joint distributions, as input and surrogate models may contain different numbers of variables. Aligned variables are those present in both input and surrogate models or within the surrogate model before and after metamodeling. Specifically, we (i) quantify model uncertainty by summing the entropies of the marginal distributions of aligned variables, (ii) compute model consistency by averaging the overlaps between these marginal distributions, and (iii) assess model complexity based on the total degree of the underlying PGM graph [47]. Other criteria, such as the number of model variables, can also be employed to measure model complexity for different representations.

### Selection of the surrogate model

A good surrogate model should reproduce the variables of interest in the input model by recapitulating their statistical dependencies, while maintaining a reasonable level of complexity. Based on the analysis of probability propagation in the conversion stage (Methods), we select the best surrogate model by enumerating and ranking surrogate models through: (i) the uncertainty of each surrogate model *i* (*ie*, the model entropy 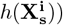; (ii) the consistency between surrogate and input models (*ie*, the average overlap index for all aligned variables 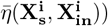;and (iii) the surrogate model complexity (*ie*, degree of the surrogate PGM graph *d*). We take the postprandial response (PR) model as an example input model [45]. This model consists of a system of ordinary differential equations (ODEs) that describes changes of insulin and glucose levels in plasma and various body tissues over time (SI Text 5, Table S1). To estimate a model distribution based on deterministic ODEs, we first assign the uncertainty using the standard error associated with the model variables, derived from the experimental data used to build the model. We construct the surrogate PGM graph (Fig. 2A) by (i) selecting eight surrogate model variables 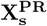 out of twelve input model variables 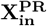 and (ii) constructing a dynamic Bayesian network (DBN) by incorporating conditional statistical dependencies derived from the ODEs as edges in the graph [48]. We compute the surrogate PGM distribution by employing probabilistic inference over the surrogate PGM graph *via* the junction tree algorithm [27].

**Fig 2.**
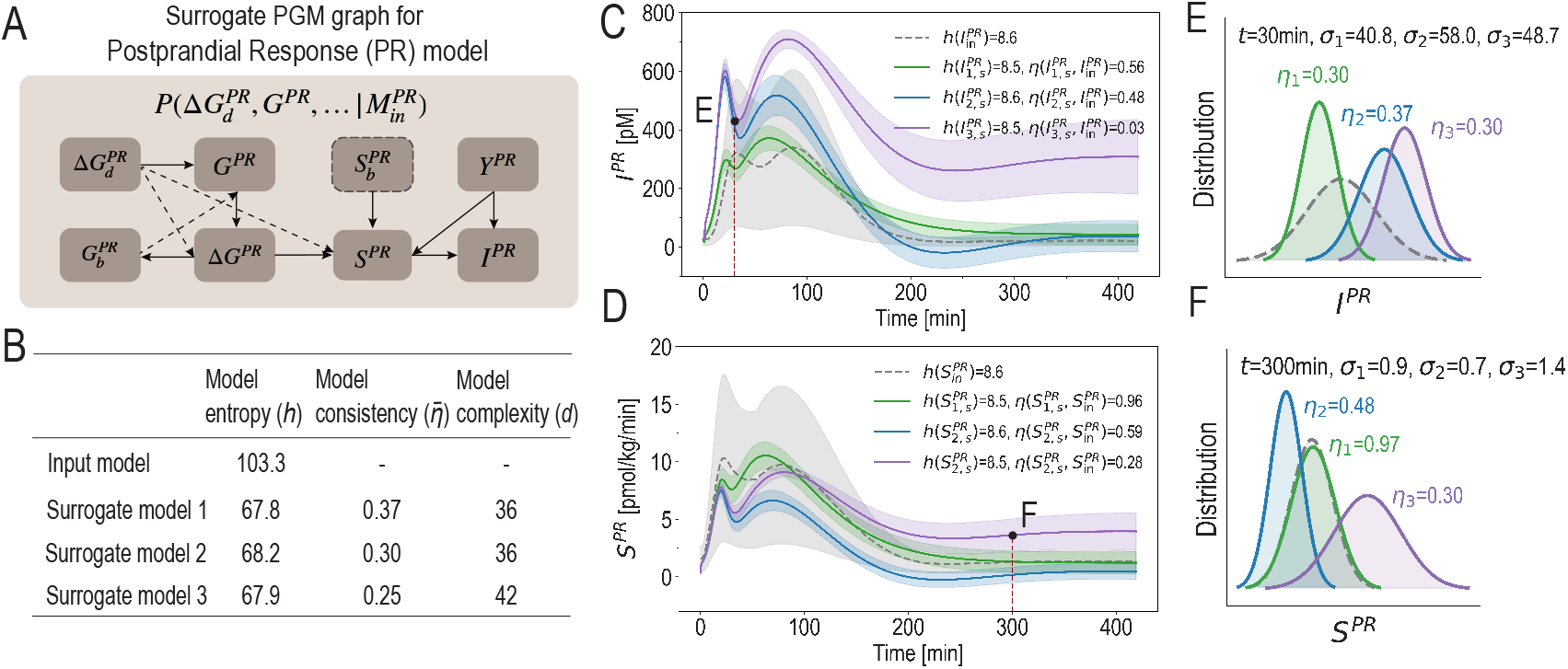
Probability propagation from input model to surrogate model. **(A)** Surrogate PGM graph for the Postprandial response (PR) input model. Frames indicate surrogate model variables, while dashed frame indicates time-independent variables, dashed edges indicate additional edges in surrogate model 3. **(B)** Table for model uncertainty 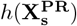,model consistency 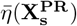 and model complexity *d* of two computed surrogate models. **(C-D)** Time courses of variables *I*^*P R*^ and *S*^*P R*^ in the input model (dashed gray line), surrogate model 1 (solid green line), surrogate model 2 (solid blue line), and surrogate model 3 (solid purple line). Dashed or solid lines represent the mean values, while shaded areas represent the standard deviations. 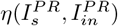 and 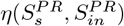 represent the average overlaps of the marginal distributions of the aligned variables in input and surrogate models over 420 min. **(E-F)** Distributions of *I*^*P R*^ and *S*^*P R*^ at 30 min. *η* denotes the overlap of the marginal distributions in input and surrogate models, while *σ* denotes the standard deviation of the variable marginal distribution.

In principle, we should enumerate the degrees of freedom in the conversion methods to select the optimal surrogate model. This involves the number of surrogate model variables, as well as all conceivable graph structures and conditional probability distributions (CPDs) in the surrogate PGM graph. To illustrate, we construct three surrogate models for the postprandial response (PR) input model with different complexities 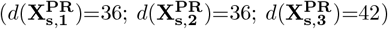 (Fig. 2A, SI Table S4). Surrogate model 1 exhibits lower entropy 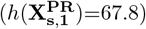 and higher consistency 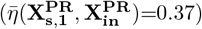 compared to surrogate model 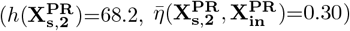; and similar model uncertainty and higher model consistency compared to surrogate model 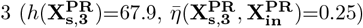 (Fig. 2B). As expected, variations in entropy and consistency of individual variables or the same variable at different time points may not necessarily align with those of the entire models (Fig. 2C-F). For example, the plasma insulin level over 420 min in surrogate model 1 shows similar entropy 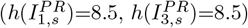 and significantly higher consistency 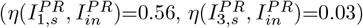 ompared to surrogate model 3 (Fig. 2C). However, the plasma insulin level at 30 min in surrogate model 1 shows a lower variable uncertainty (*σ*_1_ *< σ*_3_) and a similar overlap index (*η*_1_ = *η*_3_) compared to surrogate model 3 (Fig. 2E). Variations are also observed for the insulin secretion rate *S*^*P R*^ at different time points (Fig. 2DF). Among three candidate surrogate models, the measures of model uncertainty, consistency, and complexity guide us in selecting the optimal surrogate model, *ie*, the surrogate model 1.

### Selection of the coupling PGM graph

A good coupling PGM graph should minimize uncertainty of surrogate models after metamodeling. It should also maximize consistency among surrogate models before and after metamodeling, *ie*, changes them minimally while still achieving their statistical dependency restrained by coupling, thus rationalizing them. Based on the analysis of probability propagation in the coupling stage (Methods), we select the best coupling PGM graph by enumerating and ranking coupling PGM graphs through: (i) the overlap between the model variable distribution and its reference *η*(*X*_*s,upd*_, *X*_*ref*_), if ground truth or gold standard is available; (ii) the updated surrogate model uncertainty (*ie*, the sum of entropy for all updated surrogate models 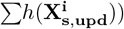;(iii) the consistency among surrogate models (*ie*, the average overlap index for all surrogate model variables before and after metamodeling 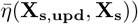;and (iv) the model complexity resulted from specific coupling PGM graphs (*ie*, degree of the meta PGM graph *d*). To ensure uniformity in selecting the best coupling PGM graphs with and without references, we introduce the coupling variables along with their observation variables (SI Table S5). The priors for these observation variables are estimated from the distributions of connecting variables in all coupling PGM graphs, or from a reference if available. Differences in model uncertainty and model consistency with and without observation variables of coupling variables, are detailed in SI Fig. S6.

We construct the coupling PGM graph by identifying six connecting variables, introducing four coupling variables, and specifying seven CPDs over a head-to-tail coupling PGM graph (Fig. 3A). First, we identify the connecting variables using: (i) model variable semantics (*eg*, both 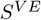 in the VE model and 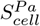 in the Pa model describe the rate of insulin secretion of a cell); (ii) prior knowledge about the statistical relations between connecting variables (*eg*, the plasma glucose concentration 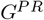 in the PR model has been estimated to be double the intracellular glucose concentration 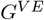 in the VE model [49]); and (iii) statistical analysis (*eg*, 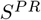 in the PR model and 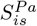 in the Pa model present a Pearson correlation coefficient of 0.94 in their corresponding surrogate models) (SI Fig. S2). Second, we introduce the coupling variables to formally relate the connecting variables by (i) introducing variables of the same semantics as the connecting variables (*eg*, the coupling variable 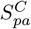 relates 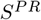 and 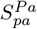,all describes the rate of insulin secretion of a pancreas); and (ii) introducing new variables to depict statistical dependencies between the connecting variables (*eg*, the coupling variables 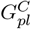 and 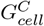 relate the glucose concentration in plasma 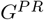 and in the cell *G*^*PR*^). Third, we specify the CPDs between the connecting variables and the coupling variable(s) using (i) statistical dependencies based on prior knowledge (*eg*, three CPDs are specified over the connecting variables 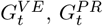,and the coupling variables 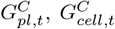 in the coupling PGM graph 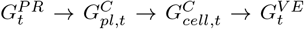 based on experimental measurements [49]):

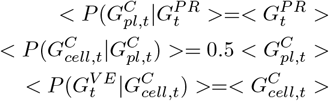

where *<>* denotes the mean of a Gaussian distribution); and (ii) exhaustive sampling (see below, Selection of the coupling PGM graph without reference). We specify CPDs across time slices in the coupling PGM graph by defining both the intra-timeslice statistical dependencies between the coupling variables and the connecting variables, and by the intra-timeslice or inter-timeslice statistical dependencies over variables in individual surrogate models. For example, CPD of the time-dependent variable 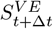 in the coupling graph is defined by the coupling variable at the same time slice 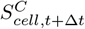,the connecting variables at the previous time slice 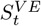 and other variables in the surrogate VE model, denoted as 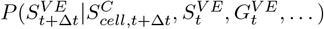 (SI Table S6).

**Fig 3.**
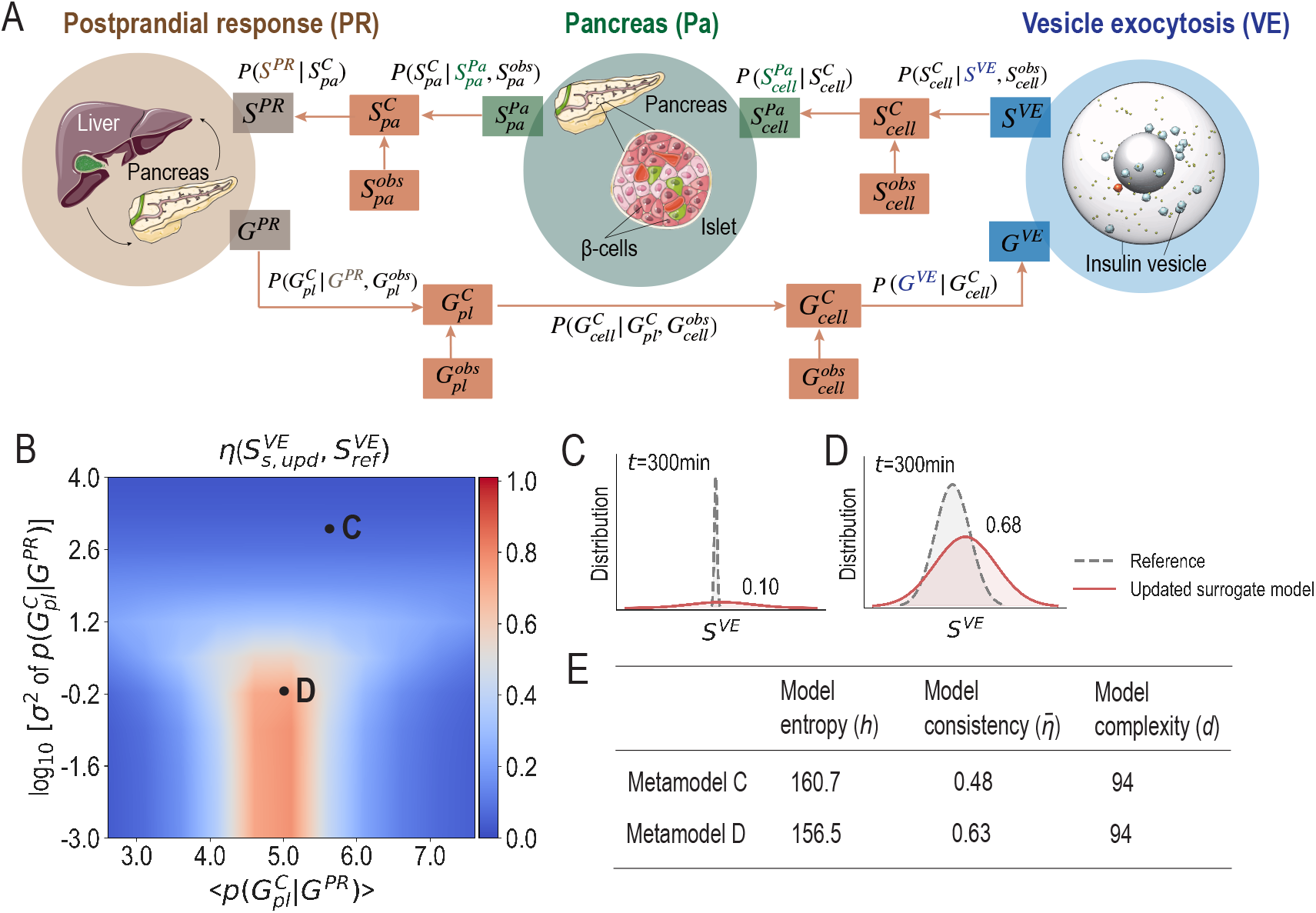
Optimization of metamodeling by improving the selection of coupling PGM graphs with reference. **(A)** Schematic illustration of the GSIS metamodel. The model PGM graphs of the Postprandial Response (PR), Pancreas (Pa), and Vesicle Exocytosis (VE) models are shown in khaki, green, and blue, respectively. The coupling PGM graphs are shown in orange. **(B)** The distribution overlaps between the updated surrogate model variable 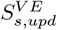 and the reference variable 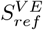 as a function of the means (x-axis) and variances (y-axis) of the 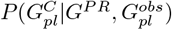 in one selected metamodel in A. **(C-D)** Comparison of the reference and updated surrogate model variable distribution at 100 min and 300 min for two different metamodels in B. **(E)** Table of model uncertainty, model consistency and model complexity of the metamodels selected in B.

### Selection of the coupling PGM graph with references

When we have access to the ground truth or the gold standard of certain model variables (*eg, S*^*V E*^), our aim is to find a coupling PGM graph that maximizes the overlap 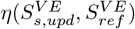 between the variable *S*^*V E*^ and its reference. To illustrate, we construct 121 metamodels with different means and variances of 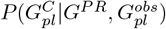 in one coupling PGM graph structure (Fig. 3A), resulting in varying 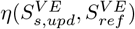 values (Fig. 3B). When the variance *σ*^2^ of 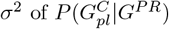 exceeds 10 mM, the overlap is seen to remain low regardless of the mean 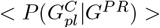 (blue region in Fig. 3B). For instance, at 300 min, the overlap between the distribution of *S*^*V E*^ and its reference is 0.1 (point C in Fig. 3B, Fig. 3C). When the variance is below 10 mM, a region of substantial overlaps emerges when sampling different means (4.5 mM - 5.5 mM, red region in Fig. 3B). The maximal overlap occurs at a mean of 5.1 mM and a variance of 1 mM (point D in Fig. 3B), with a value of 0.68 at 300 min (Fig. 3D). Thus, we find the optimal coupling PGM graph that provides the most reliable estimation for *S*^*V E*^ compared to its reference among the candidate graphs (point D in Fig. 3B). Furthermore, we observe a lower model uncertainty (*h*_*D*_ *< h*_*C*_) and a higher model consistency 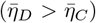 resulting from the optimized coupling PGM graph compared to the non-optimized one (Fig. 3E).

### Selection of the coupling PGM graph without references

In practice, however, we might not have access to the reference of any model variables. Our aim is to find a coupling PGM graph that minimizes the entropy of surrogate models (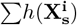 and 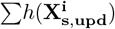) and maximizes the consistency among surrogate 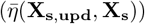,as well as reducing the complexity of the metamodel (*d*(**X**_**meta**_)). Ideally, we enumerate the degrees of freedom in the coupling strategies to select the optimal coupling PGM graph. This involves the number of connecting variables, the number of coupling variables, all conceivable graph structures, and CPDs in the coupling graph. To simplify, we showcase these degrees of freedom separately.

First, we select the connecting and coupling variables based on model variable semantics, prior knowledge, and statistical analysis, as detailed above and in SI Fig. S2. Next, we construct three coupling PGM graph structures using different sets of connecting and coupling variables (Fig. 4A-C). Note that we use CPDs listed in SI Table S6 to illustrate the selection of coupling PGM graph structures, rather than conducting an exhaustive search for the optimal set of CPDs for each candidate graph structure. Among the constructed graphs, G1 exhibits the lowest model uncertainty (*h*_1_=156.5) and the highest model consistency 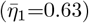 compared to the other two (Fig. 4A-C). Thus, we select the optimal coupling PGM graph structure G1, despite its relatively high model complexity (*d*_1_=94). Note that the structure and CPDs in the selected coupling PGM graph G1 (Fig. 4A) are identical to those shown in Fig. 3. Third, we construct a total of 121 metamodels with varying means and variances of 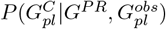 to select its optimal CPD. Noteworthy, we are guaranteed to find one with the maximal model consistency by not changing the surrogate models, *ie*, the model consistency 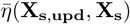 is 1, indicating that the surrogate models are not effectively coupled or rationalized. To avoid this issue, we implement restraints as follows: (i) we define distributions for the observations of all coupling variables (*eg*, 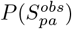 in Fig. 4A) based on the marginal distributions of two connecting variables before metamodeling (SI Fig. S3) using this equation:

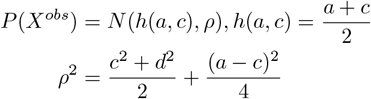

(ii) we specify the means and variances of three CPDs, 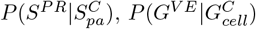 and 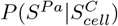 in the coupling PGM graph based on prior knowledge about their statistical dependencies. Under these restraints, we sample the CPDs introduced between the coupling variables, their observations and the connecting variables (*eg*, the mean of 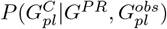 equals to 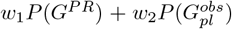 and *w*_1_ + *w*_2_=1). As shown in Fig. 4D-E, with the variance below 10 mM, the model uncertainty after metamodeling 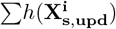 is lower than that before metamodeling 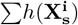 (blue region in Fig. 4D), the model consistency 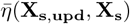 achieves a maximal region with its mean ranging between 4 mM and 6 mM (red region in Fig. 4E). Thus, among the 121 CPDs in coupling PGM graph G1, the optimal CPD for 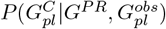 is found at a mean of 5.1 mM and a variance of 1 mM, resulting in minimal model uncertainty (point F in Fig. 4D) and maximal model consistency (point F in Fig. 4E). Another example of the selection of CPDs is conducted for 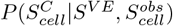 in the coupling PGM graph G1 (SI Fig. S8).

**Fig 4.**
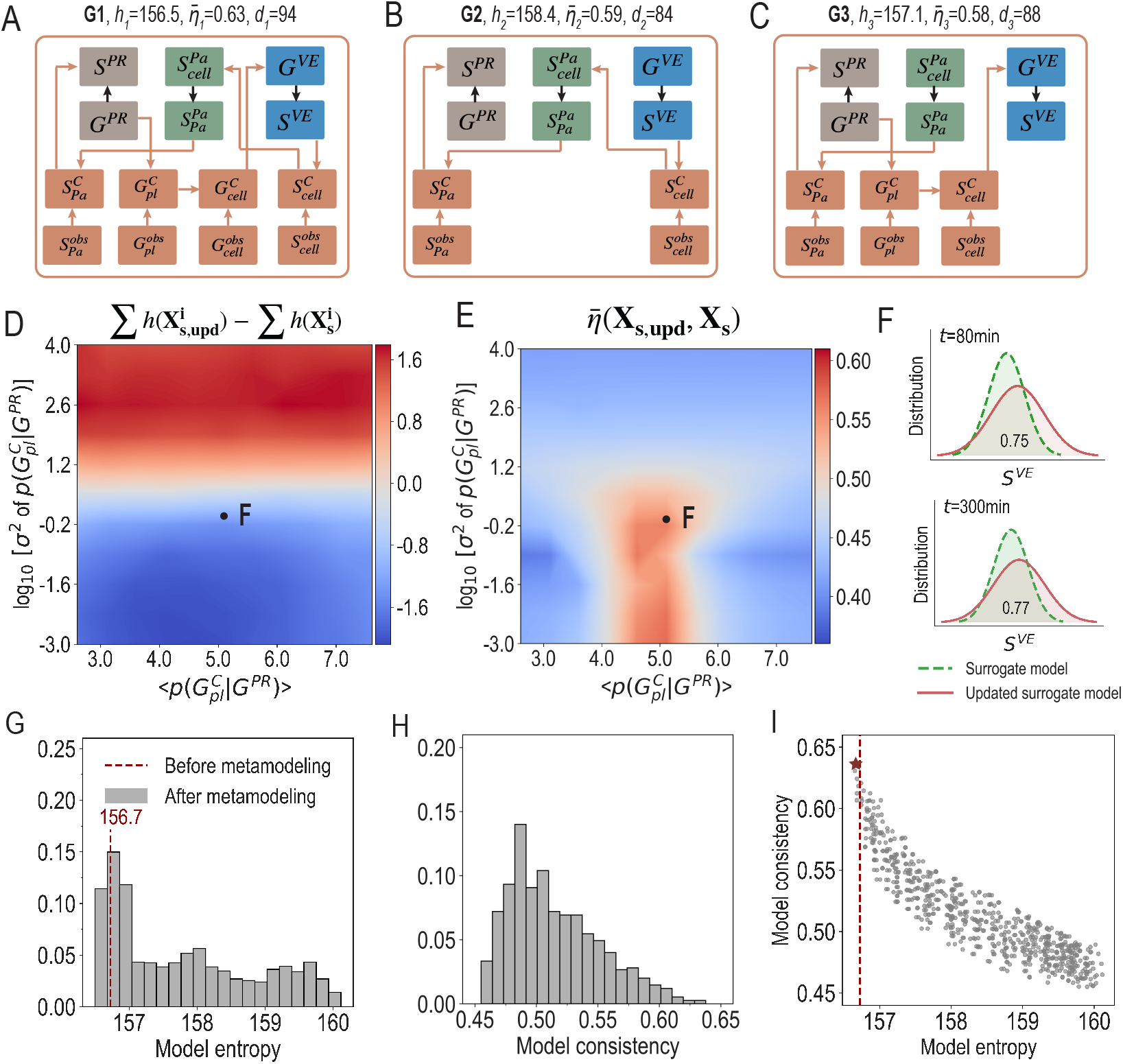
Optimization of metamodeling by improving the selection of coupling PGM graphs without reference. **(A-C)** Three coupling PGM graphs. Orange nodes and edges represent coupling variables and CPDs in the coupling PGM graphs. Model uncertainty and model consistency values are shown in figure titles. **(D)** Difference between model uncertainty before 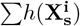 and after 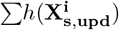 metamodeling as a function of the means (x-axis) and variances (y-axis) of the CPD 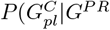 in coupling PGM graph G1. **(E)** Model consistency 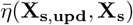 as a function of the same sets of means (x-axis) and variances (y-axis) in D. Values in D-E are computed over the metamodel timespan (420 min). **(F)** Comparison of the model variable *S*^*V E*^ distribution before and after metamodeling at 80 min and 300 min. **(G)** Distribution of model uncertainty after metamodeling with varying CPDs of 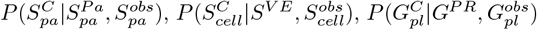, and 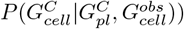.The sum of the entropy of three surrogate models before metamodeling is shown as the red dashed line. **(H)** Distribution of model consistency after metamodeling with the same set of CPDs as in G. **(I)** Change of model consistency along with the change of model uncertainty across all metamodels. The selected coupling PGM graph is shown as the red star.

In practice, one should conduct exhaustive sampling for both coupling graph structures and CPDs within each graph structure simultaneously. Upon the selected coupling PGM graph G1 (Fig. 3, Fig. 4A), we construct a total of 1,296 metamodels with varying means and variations of four CPDs, 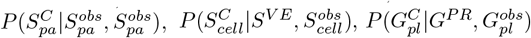, and 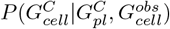.We observe that well-constructed CPDs in the coupling PGM graph can significantly minimize the overall surrogate model uncertainty (Fig. 4G). Consistency among surrogate models varies from 0.45 to 0.73 with different CPDs, indicating differences in how well the surrogate models are rationalized (Fig. 4H). Interestingly, we find an approximately negative correlation between model uncertainty and model consistency for the enumerated CPDs in the coupling PGM graph (Fig. 4I). Indeed, we wish to build a metamodel with the minimal uncertainty of the updated surrogate models and the maximal consistency among surrogate models (*ie*, star in Fig. 4I, parameter values in SI Table S6, variable plots in SI Fig. S4). Thus, among the 1,296 CPDs in coupling PGM graph G1, changes in model uncertainty and model consistency before and after metamodeling guide us in selecting the optimal CPDs for the coupling PGM graph, *ie*, star in Fig. 4I. Changes in the entropy of individual surrogate models and the standard deviations of coupling variables before and after metamodeling are detailed in SI Table S7.

### Including observations for the model system to resolve conflicts

Conflicts between surrogate models can be identified using a metamodel by comparing certain model variable distribution with its reference *η*(*X*_*s,upd*_, *X*_*ref*_) if available, model uncertainty 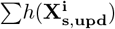 and model consistency 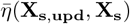.We construct 121 surrogate models with varying errors 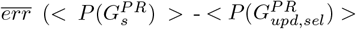, where 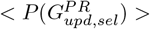 is taken from the selected optimal metamodel in Fig. 4I) and variances *σ*^2^ of 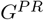.When the reference for the model variable *S*^*V E*^ is available, changes of the overlap 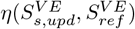 generally align with changes in the absolute error of *G*^*P R*^ (Fig. 5A). Without the reference, we observe increased model uncertainty with high variance of *G*^*P R*^, even with low errors (red region in Fig. 5B); and decreased model consistency with high error, even with low variances (blue region in Fig. 5C). Model conflicts are thus identified with relatively large errors and variances in *G*^*P R*^ (point Y in Fig. 5A-C) compared to low errors and variances (point X in Fig. 5A-C).

**Fig 5.**
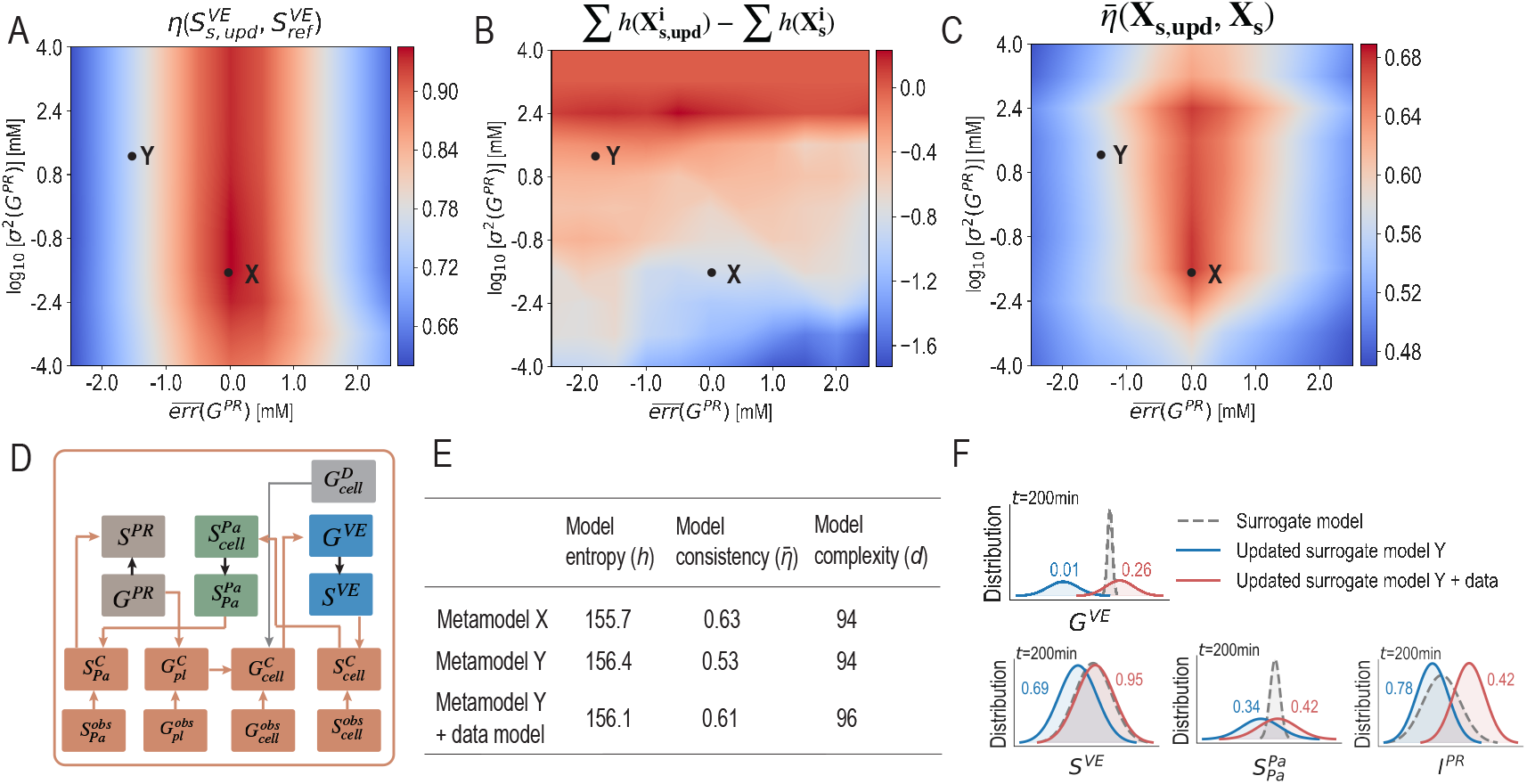
Optimization of metamodeling by solving model conflicts. **(A)** The distribution overlaps between the updated surrogate model variable 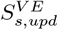 and the reference variable 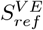 as a function of the errors (x-axis) and logarithm variances (y-axis) of the surrogate model variable *G*^*P R*^. **(B)** Difference of entropy before and after metamodeling as a function of the same sets of errors (x-axis) and logarithm variances (y-axis) in A. **(C)** Model consistency between surrogate and updated surrogate models as a function of the same sets of errors (x-axis) and logarithm variances (y-axis) in A. **(D)** Coupling PGM graph with an additional data model. **(E)** Table of model uncertainty, model consistency and model complexity of the selected metamodels (point X and point Y) in B-C and the metamodel with additional data model. **(F)** Comparison of surrogate model variable distributions at 200 min before (gray) and after metamodeling with (red) and without (blue) an additional data model.

To optimize the metamodel with conflicting surrogate models, we can (i) include additional observations of certain model variables from new experimental measurements; and (ii) remove surrogate models that are deemed poorly constructed. Here, we include an observation for the intracellular glucose concentration *G*_*cell*_ as a data model 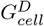 into the GSIS metamodel (Fig. 5D, SI Fig. S7). As expected, this results in an optimized metamodel, with similar model uncertainty (*h*(**X**_**s**,**upd**,**Y**_)=156.4, *h*(**X**_**s**,**upd**,**data**_)=156.1), improved model consistency 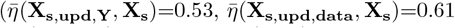, Fig. 5E), and slightly higher model complexity (*d*(**X**_**s**,**upd**,**Y**_, **X**_**s**_)=94, *d*(**X**_**s**,**upd**,**data**_, **X**_**s**_)=96) compared to the metamodel with conflicts. Moreover, we extract distributions of four variables at 200 min in different surrogate models before and after metamodeling: the intracellular glucose concentration *G*^*V E*^, the insulin secretion rate of one *β*-cell *S*^*V E*^, the pancreatic insulin secretion rate 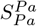,and the plasma insulin concentration *I*^*P R*^. In the optimized metamodel (red solid curve in Fig. 5F), distributions of three variables 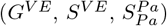 exhibit lower uncertainties after metamodeling and higher overlaps before and after metamodeling, while opposite changes are observed for *I*^*P R*^ (Fig. 5F). Till here, the metamodel is further optimized by identifying and resolving potential conflicts through the inclusion of observations of model variables as an additional data model.

### Comparison of optimized and non-optimized metamodel

Thus far, we have optimized the GSIS metamodel (star in Fig. 4I), which outperforms the previously constructed non-optimized one [1]. Specifically, the optimized PR surrogate model achieved a decrease in uncertainty 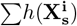 from 68.2 to 67.8, and an increase in consistency between the input and surrogate model 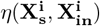 from 0.30 to 0.37, compared to the non-optimized one (Fig. 2B). The optimized metamodel achieved a decrease in surrogate model uncertainty 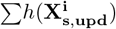 from 159.6 to 156.7, and an increase in consistency among the surrogate models 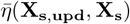 from 0.52 to 0.63, compared to the non-optimized one (Fig. 4A). We note that the GSIS metamodel can be further optimized, as model optimization is often a continuous process. For instance, this can be achieved through more comprehensive sampling of the model space using additional computational resources or through iterative refinements as new information or models become available.

With the optimized GSIS metamodel in place, we compare changes in key variables between normal and T2D subjects (Fig. 6). The mean fold change in plasma insulin level (*I*^*P R*^) predicted by the optimized metamodel (1.6) closely aligns with experimental measurements (1.7) [50], whereas the non-optimized model predicts a lower value (1.2) (Fig. 6AD). For the insulin secretion rate (*S*^*P R*^), the optimized metamodel (*eg*, 1.2±3.7 at 5 min and 2.5±2.5 at 30 min) illustrates impaired rapid insulin secretion in response to glucose stimulation more effectively than the non-optimized model (*eg*, 0.7±2.8 at 5 min and 1.2±2.3 at 30 min) (Fig. 6BD) [51]. The effective rate of insulin vesicle trafficking toward the cellular periphery 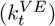 in the optimized metamodel (*eg*, 1.3±1.0 at 30 min) shows reduced trafficking efficiency in T2D subjects, consistent with total internal reflection fluorescence (TIRF) imaging measurements [52], in contrast to the increased efficiency predicted by the non-optimized model (*eg*, 0.7±1.0 at 30 min) (Fig. 6CD). Additionally, the optimized metamodel provides accurate estimates of the elevated basal plasma glucose level 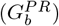 in T2D subjects (9.2±1.1 mM) compared to normal subjects (5.1±1.1 mM) (Fig. 6D) [53, 54]. These results highlight the significantly improved accuracy of the optimized GSIS metamodel in quantitatively estimating of impaired *β*-cell function in T2D subjects, which in turn underscores the critical role of optimization. In summary, our method facilitates the construction of accurate, precise, and sufficiently simple models following the Bayesian *Occam’s razor* rationale. Through continuous and iterative optimization, we enhance the reliability and applicability of these models, improving our understanding of complex biological systems such as whole cells and intricate diseases.

**Fig 6.**
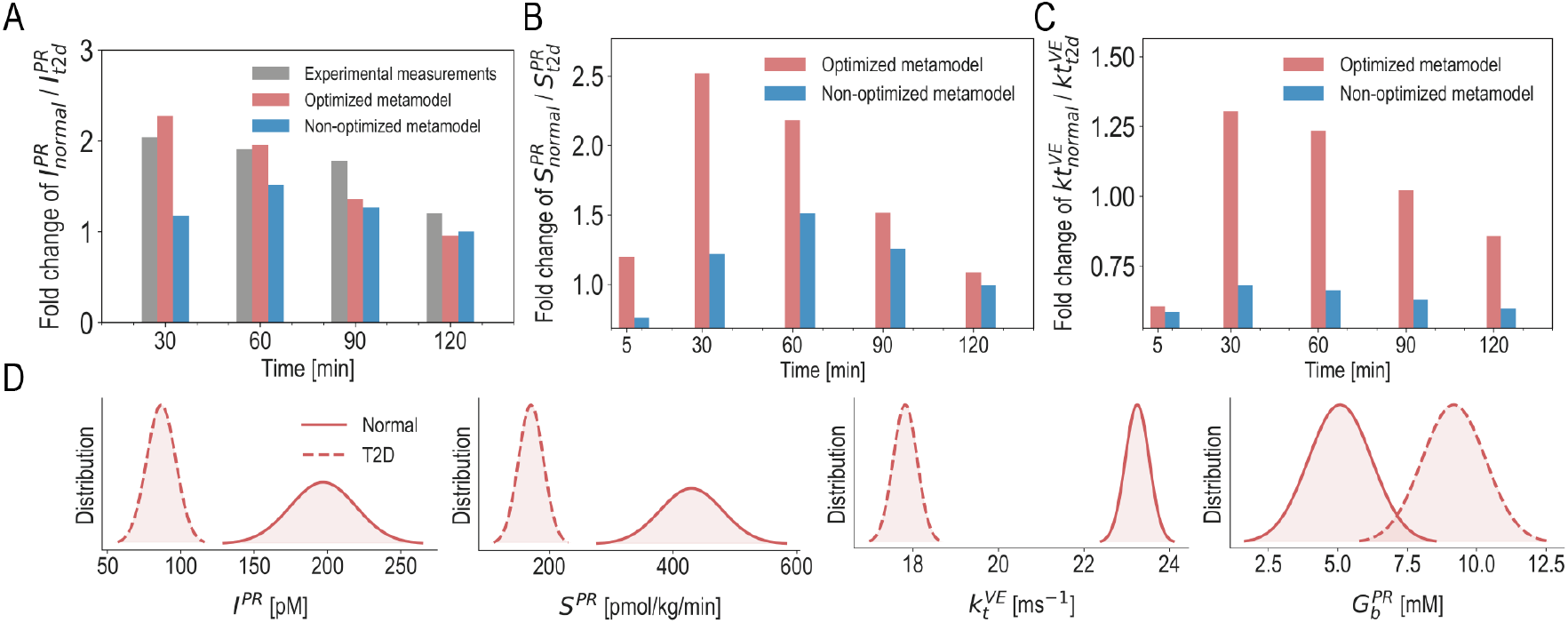
Comparison of the optimized and non-optimized metamodel. **(A-C)** Fold changes of variables *I*^*P R*^, *S*^*P R*^, and 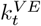 in normal and T2D subjects, as predicted by the optimized (red) and non-optimized (blue) GSIS metamodels. Experimental measurements from 36 normal subjects and 36 T2D subjects are shown in grey. **(D)** Distributions of the model variables 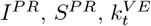,and 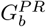 at 30 minutes in normal (solid line) and T2D (dashed line) subjects, as predicted by the optimized GSIS metamodel.

## Discussion

Modeling complex systems inherently involves high levels of complexity due to various inputs and inherent uncertainties. Integrative modeling integrates available input information, while metamodeling integrates a collection of input models that describe different aspects of the entity of interest. Taking metamodeling as our case study, we present the probability calculus using an analytical example to establish definitive rules for the probability propagation from input models to the metamodel and propose quantitative assessments for it (Fig. 1, Methods). Based on these, we have developed a method to optimize metamodeling for complex biological systems by: (i) minimizing model uncertainty, (ii) maximizing consistency between input and surrogate models to ensure the reproduction effectiveness in the conversion stage, as well as among surrogate models while changing them minimally to rationalize them in the coupling stage; and (iii) reducing the model complexity, following the Bayesian *Occam’s Razor* rationale. We showcase its benefits in the selection of surrogate models and coupling PGM graphs (Fig. 2-4), as well as including observations of model variables as an additional data model to resolve conflicts among surrogate models (Fig. 5). This method facilitates the construction of an optimized metamodel that yields more accurate estimates of impaired *β*-cell dynamics and function in T2D subjects (Fig. 6), thereby underscoring the critical role of optimization in improving model reliability and applicability.

Moreover, facilitated by our method, the iterative nature of various integrative modeling approaches provides a dynamic framework for continuous optimization. If the optimal model is not found or additional input information becomes available, we revisit the construction stages iteratively, aiming to continuously improve its precision, consistency while ensuring simplicity [55, 1]. For example, in metamodeling, the iterative selection of the surrogate model can be achieved by assessing the uncertainty, consistency, and complexity when converting the input model into a surrogate model, as well as evaluating the updated surrogate models in the backpropagation stage; the iterative selection of the coupling PGM graph can be achieved by assessing the uncertainty of surrogate models and the consistency among them in the backpropagation stage while considering that model complexity should be maintained at an acceptable level. New experiments may be conducted to reveal conflicts in the model, leading to potential adjustments in assumptions when constructing a model (*eg*, simplifying statistical dependencies among model variables). The optimized model is expected to provide new insights into the system and guide experiment design, which in turn drives further optimization.

Although this method is appealing, several aspects require further development to achieve model optimization in an automated manner. First, we ought to design standard scoring functions for the evaluation of the modeling process. This could potentially be achieved by combining the weighted sum of model uncertainty, model consistency, and model complexity, along with other criteria of the modeler’s interest (*eg*, model accuracy when the ground truth or gold standard is available) [56, 57]. Second, with a standard scoring function in place, we will be able to conduct exhaustive sampling to thoroughly search the space of alternative models and thus avoid specific local minima (*eg*, stochastic sampling [55], grid search [58]). Third, we should extend the algorithms to accommodate other types of distributions of model variables. For instance, several Bayesian filters can be incorporated to allow for inference of non-Gaussian models, including Kalman filters for linear transition functions, extended Kalman filters for nonlinear transition functions, and particle filters for nonlinear non-gaussian processes [59, 60, 61]. We anticipate that our method following the Bayesian *Occam’s razor* rationale will find applications in the construction and optimization of modeling for diverse complex systems, thereby further enhancing our interpretation and prediction capabilities.

## Data availability

No experimental data was generated as a part of this study. All experimental data used in this study was previously published and cited accordingly.

## Code availability

The software, input files, and example output files for the present work are available at https://github.com/salilab/metamodeling/tree/optimization.

## Acknowledgments

**General:** We express our gratitude to Andrej Sali for his valuable advice and fruitful discussions. We acknowledge helpful comments by Jared Sagendorf, Eli Draizen, and Neelesh Soni. We are grateful to all members of the Pancreatic ß-Cell Consortium for providing the context in which this research was performed. **Funding:** B.R. was supported by a Minerva Center grant on Cell Intelligence. C.W., J.H.Z., J.J.Z., X.H. and

L.S. were supported by ShanghaiTech University. We acknowledge the Shanghai Frontiers Science Center for Biomacromolecules and Precision Medicine at ShanghaiTech University, and the High-Performance Computing (HPC) Platform of ShanghaiTech University for computational resources. **Author contributions:** L.S. and X.H designed and supervised the paper, C.W., J.H.Z., J.J.Z performed the research and wrote the manuscript.

B.R. and J.S. supervised the project and provided inputs in the manuscript. **Competing interests:** The authors declare no competing financial interest. **Data and materials availability:** The data that support the plots within this paper and other findings of this study are available online.

