## Supplementary Information for "Bayesian *Occam’s Razor* to Optimize Metamodeling for Complex Biological Systems"

### Supplementary Text

#### 1. Definitions

Model variables can be classified into two groups. First, target variables ($R_m$) that are of primary interest and inform the answers to the questions asked of a model (*eg*, atomic coordinates in protein structure models); $M_m$ are the values of these variables. Second, nuisance variables ($R_r$) that quantify the modeling process (*eg*, variables specifying model representation, form of a scoring function, and parameters of a scoring function such as atomic isotropic temperature factors); $M_r$ are the values of these variables. The distinction between these two variable groups depends on the usage of the model (*eg*, isotropic temperature factors can be seen as either target variables or nuisance variables). The model variable space ($R$) and the model space ($M$) are thus defined as:

$$R = \{R_m, R_r\}, M = \{M_m, M_r\} \tag{1}$$

where $R_m$ denotes target variables, $R_r$ denotes nuisance variables, $M_m$ denotes a vector of values of target variables, $M_r$ denotes a vector of values of nuisance variables. Besides, we define the "truth" as $R_0$ (the true $R_m$ variables for the model target) and $M_0$ (the true values of $R_0$).

The propagation of probability through all three modeling steps (representation, scoring, and sampling) determines the posterior model density [1, 2]. Model validation does not change the posterior model density. Note that, the boundaries among representation, scoring, and sampling can be blurred/arbitrary because we include all variables in addition to $R_m$ into $R_r$. From a Bayesian view, the target posterior model density is proportional to the product of prior and likelihood as:

$$P(M_m|M_r, I) \propto P(I|R_r = M_r, R_m = M_m)P(R_r, R_m) \tag{2}$$

Here, $P(I|R_r = M_r, R_m = M_m)$ is the likelihood of all the model variable values. $P(R_r = M_r, R_m = M_m)$ is the prior distribution of all values of (i) all representational variables (representation types),(ii) all variables of each representation type, (iii) all scoring functional forms, (iv) all parameters of each scoring functional form, and (v) all sampling variables. The prior is usually assumed as uniform without input information. $P(M_m|M_r, I)$ is the posterior model density of all values of the target variables, given nuisance variables and input information. In terms of model selection, $P(R_r = M_r|I)$ is the distribution over all values of all variables in the modeling process given input information. We can rank different $M_r$ using $P(R_r = M_r|I)$ to find a best $M_r$ at the peak of $P(R_r = M_r|I)$ by integrating all $M_m$ for each $R_r$ as:

$$P(R_r = M_r|I) = \int_{M_m} [P(M_m, M_r|I)dM_m] = \int_{M_m} [P(M_r|M_m, I)P(M_m|I)dM_m] \tag{3}$$

#### 2. Probability Propagation in a general modeling process

Modeling is a process that uses the input information to produce a model of some "truth" (*ie*, model target), such as a system (*eg*, protein structure), a process (*eg*, the vibration of a bond), and a physical law that led to the system's (process) incarnation in nature (*eg*, molecular mechanical force field). We present probability propagation from input information to the posterior model density through five example modeling scenarios.

<u>**Example 1**</u> The modeling target is an atomic protein structure given its complete representation, we assume atoms exist in nature by definition. $R_0$ is the true position coordinates for each atom. $M_0$ is the value of the true position coordinates. $R_m$ is the coordinates for each particle (*eg*, atom). $R_r$ is empty since we were given a complete representation (*eg*, includes particles represented as atoms, particle interactions represented as covalent or L-J potentials where the functional form and parameters are well defined) in the input information. The target posterior model density $P(M_m|M_r, I)$ is thus the distribution over the values of all atomic coordinates of the protein.

**Example 2** The modeling target is a coarse-grained protein structure when some coarse-graining aspects are not given and need to be computed as part of modeling. $R_0$ is the true coordinates for each bead of the protein structure. $M_0$ is the value of the true coordinates for each bead. $R_m$ is the coordinates for each bead. $R_r$ includes the number of residues per bead $b$, the choice of quadratic or cubic terms for distances between beads $s$, and its parameters $\theta$ (*eg*, the force constant and mean distance for the harmonic terms between beads). Thus, the target posterior model density $P(M_m|M_r, I)$ is the distribution over the values of coordinates of all beads. $P(M_r|I)$ is the distribution of all variables in the modeling process $(b, s, \theta)$, given the input information. $P(M_r|I)$ is computed by marginalizing over $M_m$ (coordinates of all beads) from the joint distribution $P(M_m, M_r|I)$. In other words, we compute all $M_m$ using all $M_r$, followed by weighted averaging over $M_r$ for each $M_m$ by the probability $P(M_m|I)$.

**Example 3** The modeling target is the functional form and the parameter values of the potential energy terms $E$ for covalent bonds $b$ between defined atom types of a classical molecular mechanics force field (MMFF). In nature, there exist covalent bond forces between atoms (the "truth"). We aim to model this covalent bond force, based on the input information from a quantum mechanics (QM) model for the bond force. $R_0$ is the true potential energy function in the universe of statistical mechanics of atoms as particles (the "truth"), including the true functional form and its parameter variables. $M_0$ is the value of $R_0$. $R_m$ is the potential energy functional form variable (*eg*, harmonic or L-J potential) as well as variables for its parameters (*eg*, force constants and means in the case of harmonic potential energy). $R_r$ includes the scoring functional form for judging how close the model (*ie*, potential energy function) is to input information (*ie*, the QM model). An example of the scoring functional form $S$ is a least-squares scoring function for the potential energy computed with a potential energy function model as compared to the input information from the QM model:

$$S = \sum^{\text{pairs of atom types}} \omega_b ||E(b) - QM(b)||_2 \quad (4)$$

where the values of parameter variables $\omega_b$ need to be determined as part of this modeling example. The target posterior model density $P(M_m|M_r, I)$ is thus the distribution over the values of $E$ and $\omega_b$.

**Example 4** The modeling target is the internal coordinates of atoms of a peptide backbone. Input information includes the allowed Phi-Psi Ramachandran plot classes (alpha, beta, positive Phi). We assume the only information is the allowed rotamers. No information is used for the scoring function or for validation. The sampling process is to generate dipeptides by combining all rotamers of all residues exhaustively. The modeling process is: for each allowed rotamer, we compute all possible internal coordinates and possible overlaps; for each set of internal coordinates, we only position the residues with no overlaps. Thus, $R_0$ is the true internal coordinates. $M_0$ is the value of the true position. $R_m$ is the backbone dihedral angles Phi and Psi. $R_r$ includes representational (atomic positions) and sampling variables (variables involved in the sampling process, *ie*, assigning all combinations of Phi-Psi values for each residue). The target posterior model density $P(M_m|M_r, I)$ is thus the distribution over the backbone dihedral angles Phi and Psi.

**Example 5** The modeling target is the atomic coordinates of the protein complex with the lowest free energy, among all possible subunit conformations and complex configurations. The input information is a predefined free energy landscape of the complex and the prescription that we must minimize it. $R_0$ is the true atom coordinates of the complex with the lowest free energy. $R_m$ is the atom coordinates of the protein complex (*ie*, atomic internal coordinates or absolute coordinates degenerate in translation and orientation of the entire complex). $R_r$ is all other variables in addition to $R_m$, which includes different scoring functional forms and corresponding parameters. The goal of the scoring function is to minimize the free energy of the complex by sampling a set of complex conformations. A Possible scoring function form $\Delta G_b$ can be:

$$\Delta G_b = \Delta H - T\Delta S = G_c - (G_{p1} + G_{p2}) \quad (5)$$

The target posterior model density $P(M_m|M_r, I)$ is thus the distribution over the values of the atom coordinates in a protein complex.

#### 3. Probability Propagation in metamodeling

Metamodeling divides and conquers the modeling problem of complex biological systems by integrating a collection of input models into a metamodel. We present probability propagation from input models to the posterior model density across each stage of metamodeling.

**Conversion stage** In the first stage of metamodeling, we aim to convert each input model into a surrogate model. In principle, a surrogate model can be obtained by any approach for modeling statistical distributions, such as probabilistic graphical models (PGMs) and various deep generative models [3]. PGM graphs represent model variables as nodes and their conditional statistical dependencies as edges. $R_0$ is the true distribution of the input model variables. $R_m$ is a PGM consisting of the surrogate model variables and the edges between them. $R_r$ includes the nuisance variables of the conversion process, such as which subset of input model variables are used and which edge topologies are constructed for the surrogate model. $P(M_m, M_r|I)$ is the posterior surrogate model density.

**Coupling stage** In the second stage of metamodeling, we aim to find the best coupling PGM graph that results in maximal model consistency among surrogate models while changing them minimally. Input information is the statistical relationship between surrogate model variables and the maximization of model consistency. In a special case, when there is no prior knowledge about the analytical relations between different surrogate models, we use manual fitting or machine learning approaches to identify connecting variables in surrogate models and compute a function over them by specifying both the functional form and the parameters. $R_0$ is the true distribution of the coupling PGM graph (*ie*, the true system the metamodel wants to model). $R_m$ is the joint distribution of the connecting variables in the surrogate models and the coupling variable(s). $R_r$ is all variables in addition to $R_m$, including representational variables (the choice of edges in a coupling PGM graph and its parameters, the prior of the coupling variable), the functional form of the scoring function (to judge how close the model consistency is to one) and its parameters. $P(M_m, M_r|I)$ is the posterior metamodel density.

**Backpropagation stage** In the third stage of metamodeling, we aim to update the PGM distribution of the surrogate model and the distribution of the input model by backpropagation. The surrogate model is updated by either marginalizing out or conditioning on variables from other surrogate models defined by a metamodel, achieved through exact or approximate inference over the metamodel. For each surrogate model, $R_0$ is the true distribution of the updated surrogate model. $R_m$ is the distribution of updated surrogate model variables. $R_r$ is all variable identities in addition to $R_m$, including the selection of a targeted query based on the meta PGM graph (marginal or conditional). $P(M_m, M_r|I)$ is the posterior density of the updated surrogate model variables, thus serving as the target posterior model density of metamodeling. Additionally, the input model can be updated based on its relationship with the corresponding updated surrogate model if the modeler is interested in the changes of the input model. The updated input model can be any model between the input and surrogate model spaces. Constructing the updated input model is a separate modeling process determined by the modeler; we do not address the probability propagation associated with this process.

#### 4. The analytical metamodel example

We analytically propagate the probability of two surrogate models to a metamodel, and the updated surrogate models, for the case of two static input models in the main text. We assume the distribution of two surrogate models are both Gaussian distributions.

$$P(\mathbf{X_s^a}) = P(X^a_{1,s}) = N(\omega^a_{1,s}, \omega^a_{1,s}); P(\mathbf{X_s^b}) = P(X^b_{1,s}, X^b_{2,s}) \tag{6}$$

The two surrogate models are coupled by constructing a coupling PGM graph through three steps: (i) identify two connecting variables $X^a_{1,s}$ and $X^b_{1,s}$; (ii) introduce a coupling variable $Y$; (iii) specify two Gaussian conditional probability distributions (CPDs) over a head-to-tail coupling PGM graph $X^a_{1,s} \to Y \to X^b_{1,s}$ as:

$$P_Y(Y|X^a_{1,s}) = N(X^a_{1,s}, (\omega^a_{1,s})^2) \propto \exp\{-\frac{\|Y - \mu^a_{1,s}\|^2}{2(\omega^a_{1,s})^2}\} \tag{7}$$

$$P_{X^b_{1,s}}(X^b_{1,s}|Y) = N(Y, (\omega^b_{1,s})^2) \propto \exp\{-\frac{\|X^b_{1,s} - \mu_y\|^2}{2(\omega^b_{1,s})^2}\} \tag{8}$$

The meta PGM distribution is then defined as:

$$P(\mathbf{X_{meta}}) = P(\mathbf{X_s^a}, \mathbf{X_s^b}, Y) = P(X^a_{1,s}, X^b_{1,s}, X^b_{2,s}, Y) = P(X^a_{1,s})P(Y|X^a_{1,s})P(X^b_{1,s}|Y)P(X^b_{2,s}|X^b_{1,s}) \tag{9}$$

We compute the updated distributions of the two surrogate models $a$ and $b$ through marginalization over the meta PGM distribution:

$$P(\mathbf{X^a_{s,upd}}) = P(X^a_{1,s}) = \iiint P(X^a_{1,s}, X^b_{1,s}, X^b_{2,s}, Y)dYdX^b_{1,s}dX^b_{2,s} \tag{10}$$

$$P(\mathbf{X^b_{s,upd}}) = P(X^b_{1,s}, X^b_{2,s}) = \iint P(X^a_{1,s}, X^b_{1,s}, X^b_{2,s}, Y)dYdX^a_{1,s} \tag{11}$$

By definition, We first compute the joint distribution $P(\mathbf{X^a_s}, \mathbf{X^b_s})$ from $P(\mathbf{X_{meta}})$ by marginalizing out the coupling variable. The two factors $P(X^a_{1,s})$ and $P(X^b_{2,s}|X^b_{1,s})$ in Eq.4 can be obtained from the surrogate model distributions:

$$P(X^a_{1,s}) = N(\widehat{x^a_{1,s}}, (\sigma^a_{1,s})^2) \propto \exp\{-\frac{\|X^a_{1,s} - \widehat{x^a_{1,s}}\|^2}{2(\sigma^a_{1,s})^2}\} \tag{12}$$

$$P_{X^b_{2,s}}(X^b_{2,s}|X^b_{1,s}) = N(X^b_{1,s}, (\omega^b_{2,s})^2)) \propto \exp\{-\frac{\|X^b_{2,s} - X^b_{1,s}\|^2}{2(\omega^b_{2,s})^2}\} \tag{13}$$

Note that we only need to evaluate the nominator in the equation Eq.8 since the denominator is simply a normalization factor.

$$\begin{aligned}
P(\mathbf{X^a_s}, \mathbf{X^b_s}) &= \int P(\mathbf{X_{meta}})dY = \int P(X^a_{1,s})P(Y|X^a_{1,s})p(X^b_{1,s}|Y)p(X^b_{2,s}|X^b_{1,s})dY \qquad (14)\\
&= \int \frac{\exp\{-\frac{\|X^a_{1,s}-\widehat{x^a_{1,s}}\|^2}{2(\sigma^a_{1,s})^2}\}}{\sigma^a_{1,s}\sqrt{2\pi}} \times \frac{\exp\{-\frac{\|Y-X^a_{1,s}\|^2}{2(\omega^a_{1,s})^2}\}}{\omega^a_{1,s}\sqrt{2\pi}} \times \frac{\exp\{-\frac{\|X^b_{1,s}-Y\|^2}{2(\omega^b_{1,s})^2}\}}{\omega^b_{1,s}\sqrt{2\pi}} \times \frac{\exp\{-\frac{\|X^b_{2,s}-X^b_{1,s}\|^2}{2(\omega^b_{2,s})^2}\}}{\omega^b_{2,s}\sqrt{2\pi}}dY\\
&= K\int \exp\{-(\frac{\|X^a_{1,s} - \widehat{x^a_{1,s}}\|^2}{2(\sigma^a_{1,s})^2} + \frac{\|X^a_{1,s} - Y\|^2}{2(\omega^a_{1,s})^2} + \frac{\|X^b_{1,s} - Y\|^2}{2(\omega^b_{1,s})^2} + \frac{\|X^b_{2,s} - X^b_{1,s}\|^2}{2(\omega^b_{2,s})^2})\}dY\\
&= K\int \exp\{-\frac{\|Y - \mu_y\|^2}{2\omega_y^2} + C\}dY \qquad (15)
\end{aligned}$$

where $\mu_y$, $\omega_y$, and $C$ are defined as follows,

$$\mu_y = \frac{(\omega^a_{1,s})^2 X^b_{1,s} + (\omega^b_{1,s})^2 X^a_{1,s}}{(\omega^a_{1,s})^2 + (\omega^b_{1,s})^2} \tag{16}$$

$$\omega_y = \sqrt{\frac{(\omega^a_{1,s})^2(\omega^b_{1,s})^2}{(\omega^a_{1,s})^2 + (\omega^b_{1,s})^2}} \tag{17}$$

$$C = \frac{2X^a_{1,s}X^b_{1,s} - (X^a_{1,s})^2 - (X^b_{1,s})^2}{2(\omega^a_{1,s})^2 + 2(\omega^b_{1,s})^2} - \frac{(X^a_{1,s} - \widehat{x^a_{1,s}})^2}{2(\sigma^a_{1,s})^2} - \frac{(X^b_{2,s} - X^b_{1,s})^2}{2(\omega^b_{2,s})^2} \tag{18}$$

Based on the normalization property of Gaussian distributions, we come to the following result:

$$\int \frac{\exp\{-\frac{\|Y-\mu_y\|^2}{2\omega_y^2}\}}{\omega_y\sqrt{2\pi}}dy = 1 \Rightarrow \int \exp\{-\frac{\|Y - \mu_y\|^2}{2\omega_y^2}\}dY = \omega_y\sqrt{2\pi} \tag{19}$$

which simplifies the equation (15) as:

$$P(\mathbf{X^a_s}, \mathbf{X^b_s}) = K\sqrt{2\pi}\omega_y e^C \propto \exp\{\frac{2X^a_{1,s}X^b_{1,s} - (X^a_{1,s})^2 - (X^b_{1,s})^2}{2(\omega^a_{1,s})^2 + 2(\omega^b_{1,s})^2} - \frac{(X^a_{1,s} - \widehat{x^a_{1,s}})^2}{2(\sigma^a_{1,s})^2} - \frac{(X^b_{2,s} - X^b_{1,s})^2}{2(\omega^b_{2,s})^2}\} \tag{20}$$

It can be shown that the exponential part is a quadratic function with negative 2nd-order coefficients for $X^a_{1,s}$, $X^b_{2,s}$, and $X^b_{1,s}$. As such, $P(\mathbf{X^a_s}, \mathbf{X^b_s})$ is a joint Gaussian distribution and thus the marginal distribution $P(X^a_{1,s})$ is also a Gaussian, and can be derived and re-normalize the density function w.r.t $X^a_{1,s}$:

$$P(\mathbf{X^a_{s,upd}}) = P(X^a_{1,s}) \propto \exp\{-(\frac{(X^a_{1,s})^2 - 2X^b_{1,s}X^a_{1,s}}{2(\omega^a_{1,s})^2 + 2(\omega^b_{1,s})^2} + \frac{(X^a_{1,s})^2 - 2\widehat{x^a_{1,s}}X^a_{1,s}}{2(\sigma^a_{1,s})^2})\} \tag{21}$$

$$\propto \exp\{-\frac{\|X^a_{1,s} - \mu^a_{1,s,upd}\|^2}{2(\sigma^a_{1,s,upd})^2}\} \tag{22}$$

where its conditional mean and variance can be written as:

$$\mu^a_{1,s,upd} = \frac{(\sigma^a_{1,s})^2 X^b_{1,s} + (\omega^a_{1,s})^2 \widehat{x^a_{1,s}} + (\omega^b_{1,s})^2 \widehat{x^a_{1,s}}}{(\sigma^a_{1,s})^2 + (\omega^a_{1,s})^2 + (\omega^b_{1,s})^2} \tag{23}$$

$$(\sigma^a_{1,s,upd})^2 = \frac{(\sigma^a_{1,s})^2(\omega^a_{1,s})^2 + (\sigma^a_{1,s})^2(\omega^b_{1,s})^2}{(\sigma^a_{1,s})^2 + (\omega^a_{1,s})^2 + (\omega^b_{1,s})^2} \tag{24}$$

By comparing $\sigma^a_{1,s}$ and $\sigma^a_{1,s,upd}$ using their ratio, we obtain

$$\frac{(\sigma^a_{1,s,upd})^2}{(\sigma^a_{1,s})^2} = \frac{(\omega^a_{1,s})^2 + (\omega^b_{1,s})^2}{(\sigma^a_{1,s})^2 + (\omega^a_{1,s})^2 + (\omega^b_{1,s})^2} < 1 \tag{25}$$

which means $\sigma^a_{1,s,upd} < \sigma^a_{1,s}$. Thus, the uncertainty of the updated surrogate model $a$ decreases, compared with its surrogate model distribution before coupling. For surrogate model $b$, the probability calculus is similar.

### 5. Input and surrogate models

We assume that each input model is a joint probability distribution over all model variables and is independent of one another. For input models that lack a well-defined joint distribution, such as ordinary differential equations (ODEs), we compute the joint distribution by: (i) assigning the uncertainty of model variables based on experimental measurements using their standard deviations (SD); or (ii) estimating the uncertainty of model variables in the absence of experimental measurements by employing the average ratio of standard deviation (SD) values to their corresponding mean values for all model variables with available measurements.

**Postprandial response (PR) model** The postprandial response model describes insulin and glucose levels in the plasma and various body tissues as a function of time, following a glucose-rich meal, in normal and T2D subjects. The values of these variables are computed from the rate of glucose intake, using a system of ODEs for time dependent variables (Table S1) [4]:

$$\overline{G^{PR}(t+\Delta t)} = \Delta G^{PR}_d(t)\Delta t - k_1 I^{PR}(t)\Delta t + (1-k2)G^{PR}(t)\Delta t + k_3 \Delta G^{PR}_d(t)\Delta t \tag{26}$$

$$\overline{Y^{PR}(t+\Delta t)} = (1-\alpha^{PR})Y^{PR}(t) + \alpha^{PR}\beta^{PR}\Delta G^{PR}(t) \tag{27}$$

$$\overline{S^{PR}(t+\Delta t)} = Y^{PR}(t+\Delta t) + K^{PR}\Delta G^{PR}_d(t+\Delta t) + S^{PR}_b(t+\Delta t) \tag{28}$$

$$\overline{I^{PR}(t+\Delta t)} = (1-\gamma^{PR})I^{PR}(t) + k^{PR}_4 S^{PR}(t) \tag{29}$$

The list of time-dependent conditional probabilities in the surrogate PR model is derived as follows:

$$p(\Delta G^{PR}_d(t+\Delta t)|\Delta G^{PR}_d(t)) \sim N(0.0, 10^{-4} * \epsilon^{PR}) \tag{30}$$

$$p(G^{PR}_b(t+\Delta t)|G^{PR}_b(t)) \sim N(1, 0.01 * \epsilon^{PR}) \tag{31}$$

$$p(S^{PR}_b(t)) \sim N(34.0, 0.01 * \epsilon^{PR}) \tag{32}$$

$$p(G^{PR}(t+\Delta t)|\Delta G^{PR}_d(t), G^{PR}(t), I^{PR}(t)) \sim N(\overline{G^{PR}(t+\Delta t)}, 0.01 * \epsilon^{PR}) \tag{33}$$

$$p(Y^{PR}(t+\Delta t)|Y^{PR}(t), G^{PR}(t+\Delta t), G^{PR}_b(t+\Delta t)) \sim N(\overline{Y^{PR}(t+\Delta t)}, 0.01 * \epsilon^{PR}) \tag{34}$$

$$p(S^{PR}(t+\Delta t)|Y^{PR}(t+\Delta t), \Delta G_d^{PR}(t+\Delta t), S_b^{PR}(t+\Delta t)) \sim N(\overline{S^{PR}(t+\Delta t)}, 0.01 * \epsilon^{PR}) \tag{35}$$

$$p(I^{PR}(t+\Delta t)|I^{PR}(t), S^{PR}(t)) \sim N(\overline{I^{PR}(t+\Delta t)}, 0.01 * \epsilon^{PR}) \tag{36}$$

**Pancreas (Pa) model** The pancreas model is a linear model that relates the insulin secretion rate by individual cells with the insulin secretion rate by individual islets and an entire pancreas (Table S2) [5]:

$$\overline{S_{cell}^{pa}(t)} = S_{cell}^{C}(t) \tag{37}$$

$$\overline{S_{is}^{pa}(t)} = S_{cell}^{pa}(t) * N_c \tag{38}$$

$$\overline{S_{pa}^{pa}(t)} = S_{is}^{pa}(t) * N_i \tag{39}$$

The list of time-dependent conditional probabilities in the surrogate Pa model is derived as follows:

$$p(S_{cell}^{pa}(t)|S_{cell}^{C}(t)) \sim N(\overline{S_{cell}^{pa}(t)}, 10^{-20} * \epsilon^{pa}); \tag{40}$$

$$p(S_{is}^{pa}(t)|S_{cell}^{pa}(t)) \sim N(\overline{S_{is}^{pa}(t)}, 10^{-12} * \epsilon^{pa}); \tag{41}$$

$$p(S_{pa}^{pa}(t)|S_{is}^{pa}(t)) \sim N(\overline{S_{pa}^{pa}(t)}, 0.01 * \epsilon^{pa}); \tag{42}$$

**Vesicle exocytosis (VE) model** The vesicle exocytosis model describes spatiotemporal trajectories of vesicle exocytosis in pancreatic $\beta$-cells after glucose stimulation, given an initial cell configuration. The insulin secretion rates of the $\beta$-cells under different simulation conditions are recapitulated by manually fitting linear relationships to the Brownian dynamics simulations (Table S3) [6]:

$$\overline{G^{VE}(t)} = 1.0 * G_{cell}^{C}(t) \tag{43}$$

$$\overline{k_t^{VE}(t+\Delta t)} = 5.0 + \alpha^{VE} S^{VE}(t+\Delta t)/2 \tag{44}$$

$$\overline{N_v^{VE}(t)} = \beta^{VE} * S^{VE}(t+\Delta t) \tag{45}$$

$$\overline{N_{patch}^{VE}(t+\Delta t)} = 1.0 * N_{patch}^{VE}(t) \tag{46}$$

$$\overline{N_{ins}^{VE}(t+\Delta t)} = 1.0 * N_{ins}^{VE}(t) \tag{47}$$

$$\overline{S^{VE}(t)} = k_G^{VE} G^{VE}(t) + k_p^{VE} N_{patch}^{VE}(t) + k_i ns^{VE} N_{ins}^{VE}(t) + k_D^{VE} D_v^{VE}(t) + k_R^{VE} R_{cell}^{VE}(t) \tag{48}$$

The list of time-dependent conditional probabilities in the surrogate VE model is derived as follows:

$$p(G^{VE}(t)|G_{cell}^{C}(t)) \sim N(\overline{G^{VE}(t)}, 0.01 * \epsilon^{VE}) \tag{49}$$

$$p(k_t^{VE}(t+\Delta t)|k_t^{VE}(t), S^{VE}(t+\Delta t) \sim N(\overline{k_t^{VE}(t+\Delta t)}, 1 * \epsilon^{VE}) \tag{50}$$

$$p(N_v^{VE}(t)|S^{VE}(t+\Delta t)) \sim N(\overline{N_v^{VE}(t)}, 1 * \epsilon^{VE}) \tag{51}$$

$$p(N_{patch}^{VE}(t+\Delta t)|N_{patch}^{VE}(t)) \sim N(\overline{N_{patch}^{VE}(t+\Delta t)}, 0.01 * \epsilon^{VE}) \tag{52}$$

$$p(N_{ins}^{VE}(t+\Delta t)|N_{ins}^{VE}(t)) \sim N(\overline{N_{ins}^{VE}(t+\Delta t)}, 10^{-14} * \epsilon^{VE}) \tag{53}$$

$$p(S^{VE}(t)|G^{VE}(t), N_{patch}^{VE}(t), N_{ins}^{VE}(t), D_v^{VE}(t), R_{cell}^{VE}(t)) \sim N(\overline{S^{VE}(t)}, 10^{-20} * \epsilon^{VE}) \tag{54}$$

$$p(R_{cell}^{VE}(t)) \sim N(6, 0.01 * \epsilon^{VE}) \tag{55}$$

$$p(D_v^{VE}(t)) \sim N(0.0032, 10^{-8} * \epsilon^{VE}) \tag{56}$$

### Fig. S1.

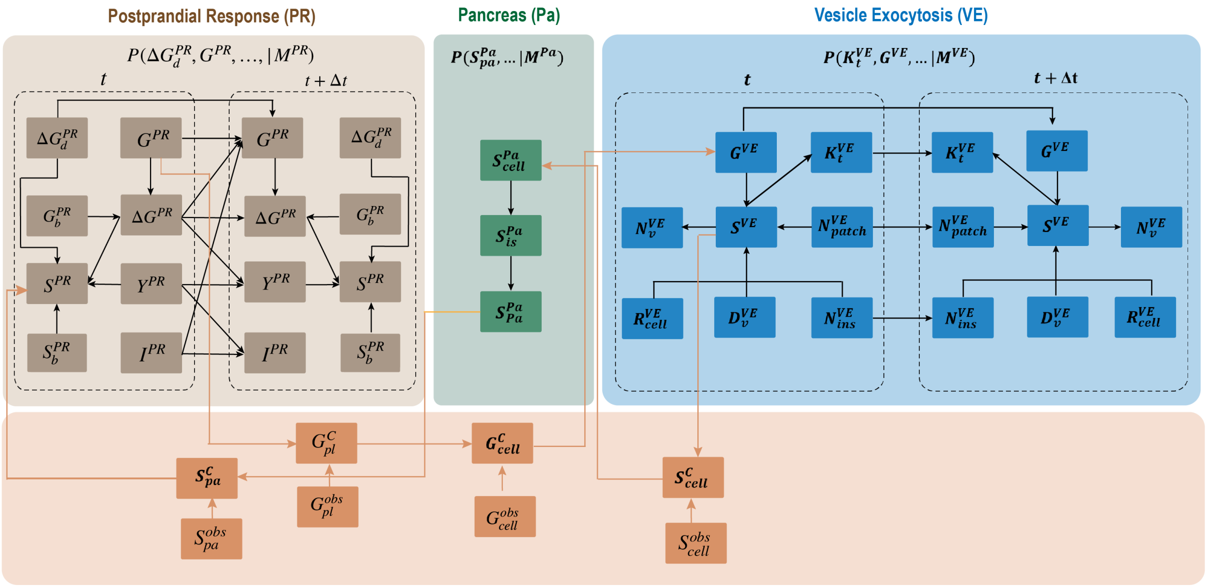


Figure 1: The meta PGM graph for the GSIS metamodel. Nodes indicate variables and directed edges indicate statistical relations between a parent and child variable in a Bayesian network; a child variable is conditionally independent of any of its non-descendants, given the values of its parent variables. The model PGM graphs of the Postprandial Response (PR), Pancreas (Pa), and Vesicle Exocytosis (VE) models are shown in khaki, green, and blue, respectively. The coupling PGM graphs are shown in orange. Superscripts indicate the model identifier. Dashed circles indicate different time slices.

#### Fig. S2.

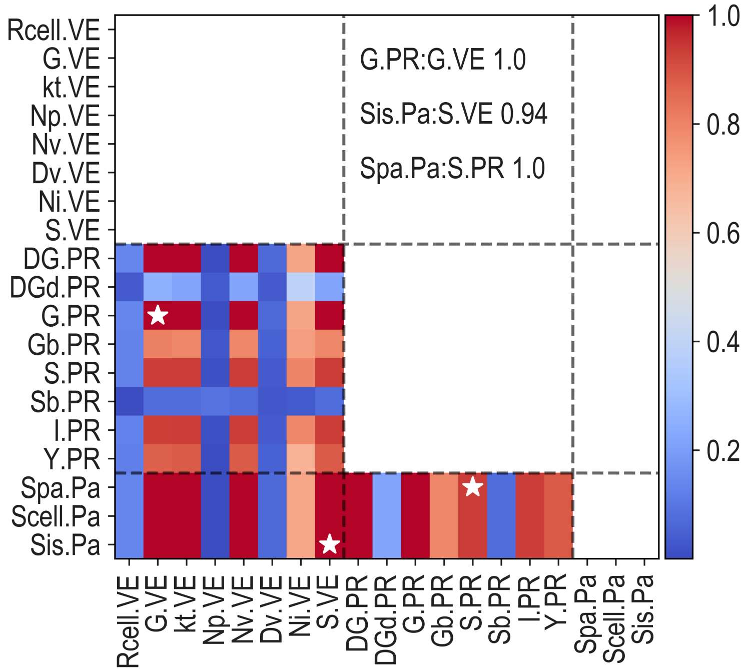


Figure 2: Identification of connecting variables through variable correlations. Correlation scores between all variable pairs from three surrogate models are computed as the average Pearson correlation coefficient over the model timespan. Three pairs of connecting variables, labeled as stars, are selected to construct the coupling PGM graph in Fig.S1.

#### Fig. S3.

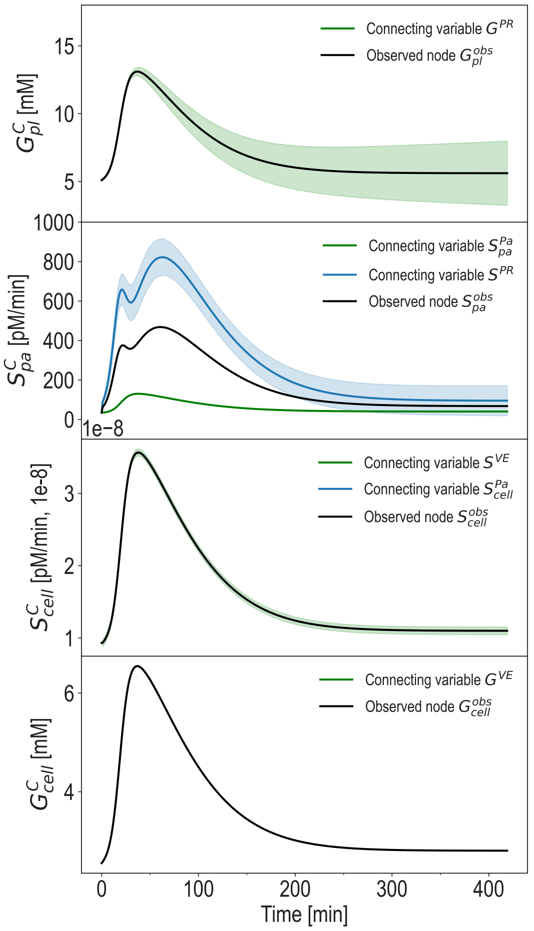


Figure 3: Time courses of the coupling variables and their observations in the coupling PGM graph in Fig.S1. The distributions of the observations are computed using Eq.39. Solid lines represent the mean values, while shaded areas represent the standard deviations.

#### Fig. S4.

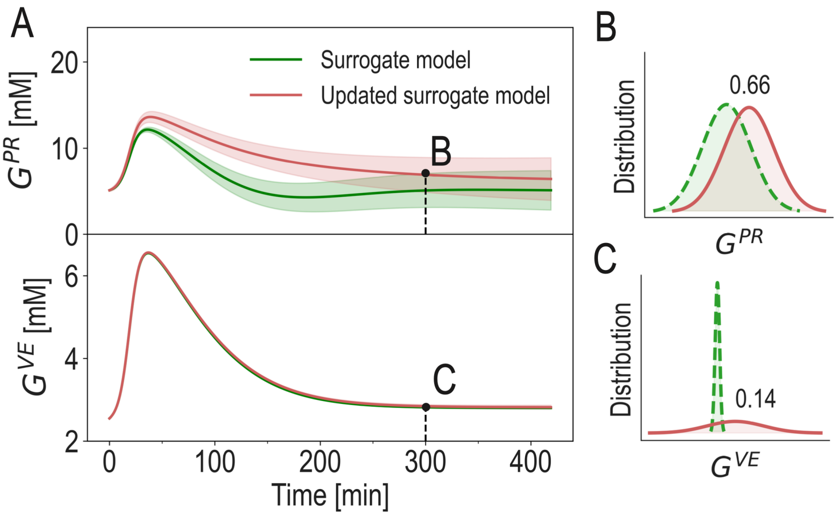


Figure 4: (A) Time courses of two selected connecting variables before and after metamodeling. Solid lines represent the mean values, while shaded areas represent the standard deviations. (B-C) Distributions of two selected model variables at 300 min.

**Fig. S5.**

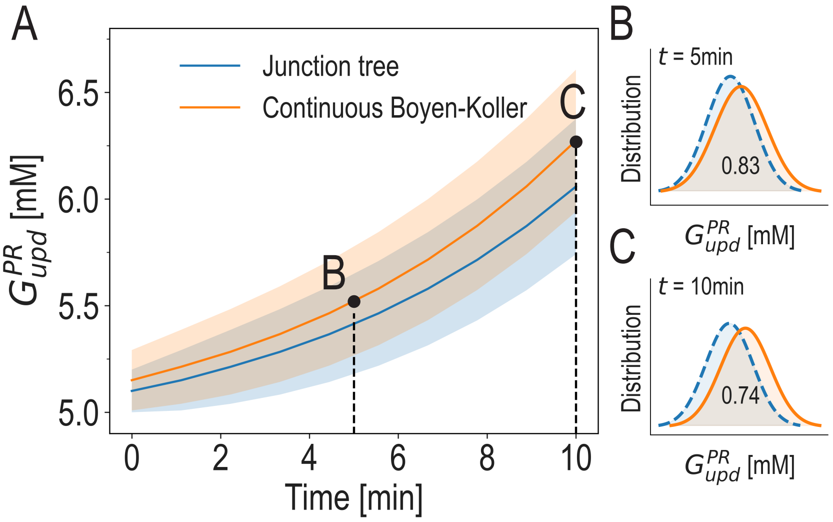


Figure 5: Comparison of different model inference methods. (A) Time course of $G_{upd}^{PR}$ computed from the optimal metamodel (star in Fig.4I in the main text) using two different inference methods: the junction tree algorithm (blue) and the continuous Boyen-Koller algorithm (orange). Solid lines represent the mean values, while shaded areas represent the standard deviations. (B-C) Distributions of $G_{upd}^{PR}$ at 5 min and 10 min.

#### Fig. S6.

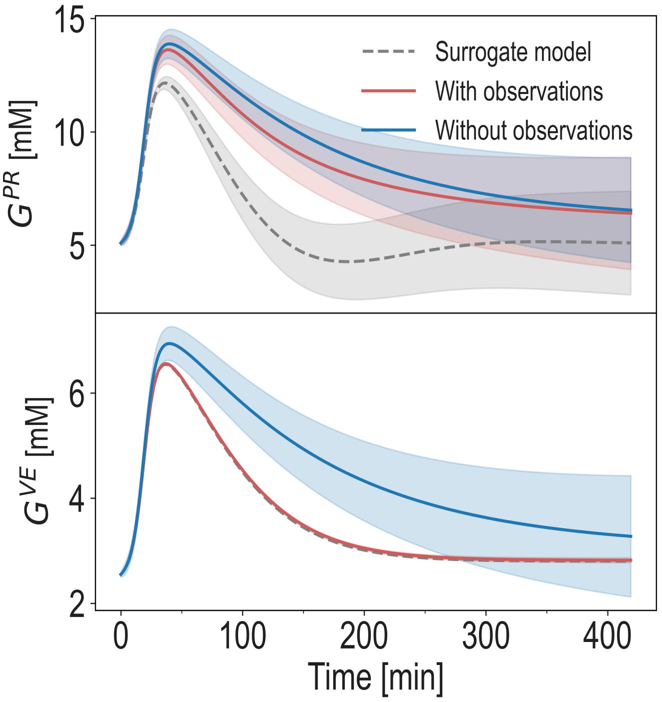


Figure 6: Comparison of metamodels with and without observations for the coupling variables. Time courses of two selected variables before metamodeling (dashed line in grey), and after metamodeling with (solid line in red) and without (solid line in blue) observations for the coupling variables. Dashed and solid lines represent the mean values, while shaded areas represent the standard deviations.

**Fig. S7.**

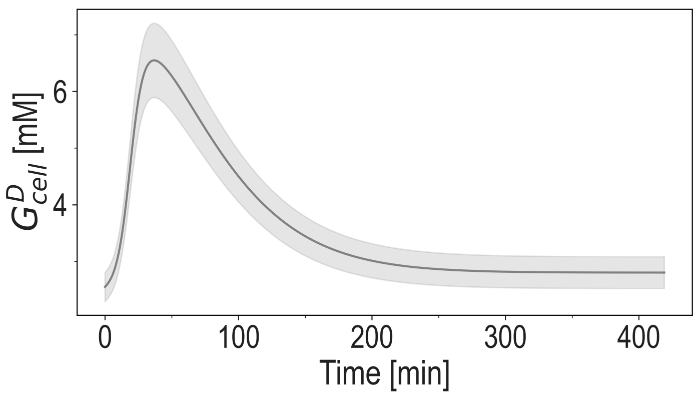


Figure 7: The data model $G^{D}_{cell}$ introduced to solve model conflicts in Fig.5C Point Y in the main text.

#### Fig. S8.

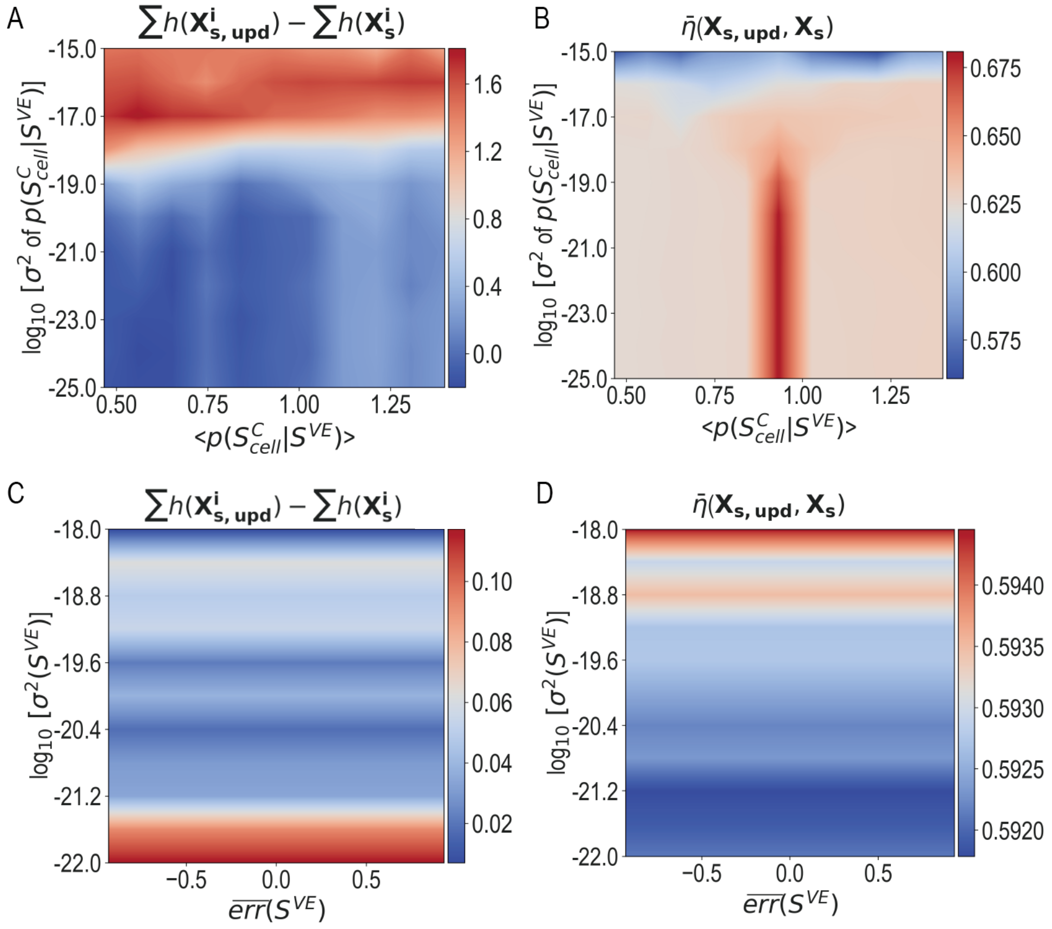


Figure 8: Comparison of model entropy and consistency by varying surrogate model variables and CPDs in the coupling PGM graph G1 in Fig.4A in the main text. (A) Change of model entropy before and after metamodeling as a function of the mean values (x axis) and variances (y axis) of 121 CPDs $P(S^C_{cell}|S^{VE})$. (B) Model consistency between surrogate and updated surrogate model distributions as a function of the mean values (x axis) and variances (y axis) of 121 CPDs $P(S^C_{cell}|S^{VE})$. (C) Change of model entropy before and after metamodeling as a function of the mean values (x axis) and variances (y axis) of the 121 marginal distributions $P(S^{VE})$ in the surrogate VE model. (D) Model consistency between surrogate and updated surrogate model distributions as a function of the mean values (x axis) and variances (y axis) of 121 marginal distributions $P(S^{VE})$ in the surrogate VE model.

### Table S1.

Surrogate model variables and CPDs in the Postprandial Response (PR) surrogate model.

| Time-dependent variables and their distributions: | | | | |
|---|---|---|---|---|
| Name | Description | Mean | Std | Unit |
| $\Delta G_d^{PR}$ | Rate of glucose intake from food digestion | 0 | 1E-2 | mM min$^{-1}$ |
| $G_b^{PR}$ | Basal plasma glucose concentration above which the cells initiate to produce new insulin | 5.1 | 1 | mM |
| $G^{PR}$ | Plasma glucose concentration | 5.1 | 0.1 | mM |
| $\Delta G^{PR}$ | Excess plasma glucose concentration compared with the basal concentration | 0 | 0.1 | mM |
| $Y^{PR}$ | Provision of new insulin to the $\beta$-cells | 0 | 0.1 | pM min$^{-1}$ |
| $S^{PR}$ | Pancreatic insulin secretion rate | 34 | 0.1 | pM min$^{-1}$ |
| $I^{PR}$ | Plasma insulin concentration | 25 | 0.1 | pM |
| $S_b^{PR}$ | Basal pancreatic insulin secretion rate | 34 | 0.1 | pM min$^{-1}$ |

### Table S2.

Surrogate model variables and CPDs in the Pancreas (Pa) surrogate model.

| Time-independent variables and their distributions: | | | | |
|---|---|---|---|---|
| Name | Description | Mean | Std | Unit |
| $S_{cell}^{pa}$ | Insulin secretion rate of a cell | 9.32E-9 | $10^{-10}$ | pM min$^{-1}$ |
| $S_{is}^{pa}$ | Insulin secretion rate of an islet | 1.0625E-5 | $10^{-6}$ | pM min$^{-1}$ |
| $S_{pa}^{pa}$ | Insulin secretion rate of a pancreas | 34 | 0.1 | pM min$^{-1}$ |
| Parameters of CPDs: | | | | |
| $N_c$ | Number of *beta*-cells in an islet | 1140 | - | - |
| $N_i$ | Number of islets in a pancreas | 3.2E6 | - | - |
| $\epsilon^{pa}$ | Variance scale of the pancreas model | 1E-3 | - | - |

### Table S3.

Surrogate model variables and CPDs in the Vesicle Exocytosis (VE) surrogate model.

| Time-dependent variables and their distributions: | | | | |
|---|---|---|---|---|
| Name | Description | Mean | Std | Unit |
| $G^{VE}$ | Intracellular glucose concentration | 2.55 | 0.1 | mM |
| $k_t^{VE}$ | Effective rate of vesicle trafficking towards the cellular periphery | 10 | 3 | $ms^{-1}$ |
| $N_v^{VE}$ | Number of insulin vesicles in one $\beta$-cell | 300 | 1 | - |
| $N_{patch}^{VE}$ | Number of activation patches per vesicle | 6 | 0.1 | - |
| $N_{ins}^{VE}$ | Amount of insulin in one vesicle | 1.8E-6 | 1E-7 | pmol |
| $S^{VE}$ | Insulin secretion rate of one $\beta$-cell | 9.32E-9 | 1E-10 | pM $min^{-1}$ |
| Time-independent variables : | | | | |
| $R_{cell}^{VE}$ | Radius of the $\beta$-cell | 6 | 0.1 | $\mu$m |
| $D_v^{VE}$ | Diffusion coefficient of insulin vesicles in the beta cell | 3.2E-3 | 1E-4 | $A^2fs^{-1}$ |
| Parameters of CPDs: | | | | |
| $\alpha^{VE}$ | Correlation between insulin secretion rate and the force coefficient of vesicle transport | 1.073E9 | - | $ms^{-1}$ $pM^{-1}$min |
| $\beta^{VE}$ | Correlation between insulin secretion rate and the number of insulin vesicles in the cell | 3.219E10 | - | $pM^{-1}$min |
| $k_p^{VE}$ | Coefficient for the number of activation patches on the vesicle surface accelerating insulin secretion rate | 8.45E-11 | - | pM $min^{-1}$ |
| $k_G^{VE}$ | Coefficient for the intracellular glucose concentration stimulating the insulin secretion | 6.59E-9 | - | pM $min^{-1}$ $mM^{-1}$ |
| $k_{ins}^{VE}$ | Coefficient for the number of insulin molecules in each vesicle determining insulin secretion rate | 0.004 | - | pM $min^{-1}$ |
| $k_D^{VE}$ | Coefficient for the vesicle diffusion promoting insulin secretion rate | 1.28E-7 | - | pM $min^{-1}A^{-2}$fs |
| $k_R^{VE}$ | Coefficient for the cell radius reducing insulin secretion rate | -2.6E-9 | - | pM $min^{-1}$ $um^{-1}$ |
| $\epsilon^{VE}$ | Variance scale of vesicle exocytosis model | 1E-3 | - | - |

### Table S4.

Different parameter values in postprandial response surrogate model 1 and 2.

Parameter values in surrogate model 1

| Name | Description | Mean | Std | Unit |
|---|---|---|---|---|
| $\alpha^{PR}$ | Delay between the glucose signal and insulin secretion | 0.05 | - | $\text{min}^{-1}$ |
| $\beta^{PR}$ | Pancreatic responsivity to glucose | 120 | - | pM $\text{min}^{-1}$ $\text{mM}^{-1}$ |
| $\gamma^{PR}$ | Transfer rate between portal vein and liver | 1 | - | $\text{min}^{-1}$ |
| $k_1^{PR}$ | Coefficient for insulin reducing glucose concentration | 0.00025 | - | $\text{min}^{-1}$ |
| $k_2^{PR}$ | Coefficient for glucose reducing glucose concentration | -0.002 | - | $\text{min}^{-1}$ |
| $k_3^{PR}$ | Coefficient for elevated glucose reducing glucose concentration | -0.001 | - | - |
| $k_4^{PR}$ | Coefficient for insulin secretion accounting for the insulin degradation | 0.4518 | - | - |
| $K^{PR}$ | Pancreatic responsitivity to the glucose rate of change | 1000 | - | $\text{pM}^{-1}$ $\text{mM}^{-1}$ |
| $\epsilon^{PR}$ | Variance scale of postprandial response model | 1E-2 | - | - |

Parameter values in surrogate model 2

| Name | Description | Mean | Std | Unit |
|---|---|---|---|---|
| $\alpha^{PR}$ | Delay between the glucose signal and insulin secretion | 0.03 | - | $\text{min}^{-1}$ |
| $\beta^{PR}$ | Pancreatic responsivity to glucose | 100 | - | pM $\text{min}^{-1}$ $\text{mM}^{-1}$ |
| $\gamma^{PR}$ | Transfer rate between portal vein and liver | 0.8 | - | $\text{min}^{-1}$ |
| $k_1^{PR}$ | Coefficient for insulin reducing glucose concentration | 0.0002 | - | $\text{min}^{-1}$ |
| $k_2^{PR}$ | Coefficient for glucose reducing glucose concentration | -0.00124 | - | $\text{min}^{-1}$ |
| $k_3^{PR}$ | Coefficient for elevated glucose reducing glucose concentration | -0.00001 | - | - |
| $k_4^{PR}$ | Coefficient for insulin secretion accounting for the insulin degradation | 0.8 | - | - |
| $K^{PR}$ | Pancreatic responsitivity to the glucose rate of change | 1000 | - | $\text{pM}^{-1}$ $\text{mM}^{-1}$ |
| $\epsilon^{PR}$ | Variance scale of postprandial response model | 1E-2 | - | - |

### Table S5.

Initial values of the observations for the four coupling variables.

| Variable | Description | Mean | Std | Unit |
|---|---|---|---|---|
| $S_{pa}^{obs}$ | Observation of insulin secretion rate of the pancreas | 34.0 | 0.005 | pM min$^{-1}$ |
| $S_{cell}^{obs}$ | Observation of insulin secretion rate of one $\beta$-cell | 9.32E-9 | 1E-20 | pM min$^{-1}$ |
| $G_{pl}^{obs}$ | Observation of plasma glucose concentration | 5.1 | 0.005 | mM |
| $G_{cell}^{obs}$ | Observation of intracellular glucose concentration | 2.5499 | 0.002 | mM |

### Table S6.

Conditional probabilities for the introduced edges in the coupling PGM graph.

| CPD | Coupling variable | [illegible] |
|---|---|---|
| $P(G^{C}_{pl,t} \| G^{PR}_{t}, G^{obs}_{pl,t})$ | $G^{C}_{pl}$ | [illegible] |
| $P(G^{C}_{cell,t} \| G^{C}_{pl,t}, G^{obs}_{cell,t})$ | $G^{C}_{cell}$ | [illegible] |
| $P(S^{C}_{cell,t} \| S^{VE}_{t}, S^{obs}_{cell,t})$ | $S^{C}_{cell}$ | [illegible] |
| $P(S^{C}_{pa,t} \| S^{Pa}_{pa,t}, S^{obs}_{pa,t})$ | $S^{C}_{pa}$ | [illegible] |
| $P(G^{VE}_{t} \| G^{C}_{cell,t})$ | - | [illegible] |
| $P(S^{Pa}_{cell,t} \| S^{C}_{cell,t})$ | - | [illegible] |
| $P(S^{PR}_{t} \| S^{C}_{pa,t})$ | - | [illegible] |

### Table S7.

Entropy of different queries before and after metamodeling.

| Query | Entropy before metamodeling | Entropy after metamodeling |
|---|---|---|
| $\mathbf{X_{meta}}$ | 202.7 | 188.6 |
| $G^C_{cell}$ | 9.0 | 9.2 |
| $S^C_{pa}$ | 9.6 | 9.0 |
| $G^C_{pl}$ | 15.6 | 8.1 |
| $S^C_{cell}$ | 11.8 | 5.8 |
| All models | 156.7 | 156.5 |
| PR model | 68.1 | 67.8 |
| Pa model | 23.9 | 24.0 |
| VE model | 64.7 | 64.7 |
